# Diffusion Tensor Imaging connectivity analysis: detecting structural alterations and their underlying substrates for Optic Ataxia in correlations with “How” stream Visual Pathways

**DOI:** 10.1101/520635

**Authors:** Ganesh Elumalai, Panchanan Maiti, Geethanjali Vinodhanand, Valencia Lasandra Camoya Brown, Nitya Akarsha Surya Venkata Ghanta, Venkata Hari Krishna Kurra

## Abstract

Optic ataxia is a neurological condition that shows clinical manifestations of disturbances in visual guided hand movements on reaching for a target object. Previous studies failed to provide substantial evidences for the neural structural pathway damaged by this condition. Therefore, this study was aimed to identify the neural structural connectivity between “Visual cortex with Superior Parietal Lobule” and to correlate its functional importance, using “Diffusion Imaging fiber Tractography. The fibers were traced, and we confirmed its extension from “Visual cortex (Brodmann’s Areas 18 and 19) to Superior Parietal Lobule (Brodmann’s Area 7)”. This new observation gives an insight to understand the structural existence and functional correlations between “Visual cortex with Superior Parietal Lobule” which is involved in targeting the grasping hand movements towards a visually perceived object, called visuo-motor coordination pathway or “how” stream pathways in visual perception. The observational analysis used thirty-two healthy adults, ultra-high b-value, diffusion MRI datasets from an Open access research platform. The datasets range between 20–49 years, in both sexes, with mean age of 31.1 years. The confirmatory observational analysis process includes, datasets acquisition, pre-processing, processing, reconstruction, fiber tractography and analysis using software tools. All the datasets confirmed that the fiber structural extension between, Visual cortex to superior parietal lobe in both the sexes may be responsible for the visual spatial recognition of objects. These new fiber connectivity evidences justify the structural relevance of visual spatial recognition impairments, such as optic ataxia.

## 1. Introduction

Many past references have proposed that the Dorsal stream visual pathway contributes to the visual-spatial recognition of an object hence, the orientation of an object in the space is achieved by the dorsal stream visual pathway. Mishkin and Ungerleider had found two streams visual pathway where the dorsal and the ventral streams contribute to locate and perceive different kinds of objects that are presented to a person, therefore, dorsal stream visual pathway is a multi-synaptic cortico-cortical pathway [1] that enables a person to guide the actions visually towards an object. Dorsal stream is an occipitoparietal pathway where there is white matter connection from the primary visual cortex (Brodmann area 17,18,19) [2] to the superior parietal lobe (Brodmann area 7). Rizzolatti and Matelli have stated that the dorsal stream visual pathway is anatomically formed by two different functional systems namely the dorso-dorsal and the ventro-dorsal streams [3]. Both the functions of the dorso-dorsal and ventro-dorsal streams are to link actions with the visual perception [3]. To grasp and to have a grip of an object is definite with the dorsal stream (occipitoparietal stream) as proposed by Culham and Stacey [4]. Milner and Goodale stated that the information provided by the dorsal stream is viewer - centered information where the object is viewer - centered, and the information is grasped [5]. Polanen and Davare mentioned that the ventral stream sends information about the object’s identity to the dorsal stream so that the dorsal stream can help to locate the object, hence we conclude by saying that firstly, the “WHAT” pathway gets activated and then the “WHERE” pathway [6], [7]. Some references mention that there is a direct white matter connection between the dorsal and ventral stream. Bilateral damage to the occipitoparietal connection can lead to optic ataxia. As our previous references haven’t mentioned any structural connection, we Team NeurON have observed a new finding that there is an existence of a neural structural connection between the primary visual cortex and the superior parietal lobe. Optic ataxia is a high order neurological deficit where the person cannot locate objects that are spatially oriented, hence, we are stating that due to the damage to the interconnection between the visual cortex and superior parietal lobe, there might be deficits in normal spatial orientation of objects. Scientists have given proof that the optic ataxia can lead to a damage to “HOW” pathway. In the case of Alzheimer’s, we found that patients present with optic ataxia which is one of the most important symptoms found in these patients of recent. Patients of Alzheimer’s with optic ataxia do not have coordination in object-oriented hand movements nor in grasping of objects [8].

## 2. MATERIALS AND METHODS

The present study design used the open access, ultra-high b-value, diffusion imaging datasets from Massachusetts General Hospital – US Consortium Human Connectome Project (MGH-USC HCP) acquired from Thirty-five healthy adults’ participants (16 Males and 16 Females, between the 20–59 years). The imaging data is available to those who register and agree to the Open Access Data Use Terms.

### 2.1 Participants

Data was acquired from thirty-two healthy adults between the ages of 20 to 59. The participants gave written consent, and the procedures were carried out in accordance with the Institutional review board (MGH/HST) approval and procedures. All participants who participated in the MGH-USC Adult Diffusion Dataset were scanned on the 3T CONNECTOM MRI scanner (see (Setsompop et al., 2013) for an overview) housed at the Athinoula A. Martinos Center for Biomedical Imaging at MGH. A custom-made 64-channel phased array head coil was used for signal reception (Keil et al., 2013).

Thirty-two healthy adults participated in this study (16 Females, 16 males, 20–59 years old; mean age =31.1 years old). All participants gave written informed consent, and the experiments were carried out with approval from the institutional review board of Partners Healthcare. Participants’ gender and age are available in the data sharing repository, Due to the limited sample size there are some ages for which we had only one participant. Given de-identification considerations, age information is provided in 5-year age bins.

### 2.2 Data Acquisition

The data was collected on the customized Siemens 3T Connectome scanner, which is a modified 3T Skyra system (MAGNETOM Skyra Siemens Healthcare), housed at the MGH/HST Athinoula A. Martinos Center for Biomedical Imaging. A 64-channel, tight-fitting brain array coil was used for data acquisition. dMRI data were acquired using a mono-polar Stejskal-Tanner pulsed gradient spin-echo planar imaging (EPI) sequence with Parallel imaging using Generalized Auto calibrating Partially Parallel Acquisition (GRAPPA). The Fast Low-angle Excitation Echo-planar Technique (FLEET) (Polimeni et al., 2015) was used for Auto-Calibration Signal (ACS) acquisitions to reduce motion sensitivity of the training data and improve stability and SNR of the GRAPPA reconstructions. The Simultaneous Multi-Slice (SMS; multi-band) technique (Feinberg et al., 2010; Feinberg and Setsompop, 2013; Setsompop et al., 2012a; Setsompop et al., 2012b) has been shown to increase the time efficiency of diffusion imaging and was considered for this protocol, however at the point in the project timeline when data acquisition was to begin, the SMS method was not implemented in the EPI sequence featuring FLEET-ACS for GRAPPA. Because of the key benefits of in-plane acceleration for improved EPI data quality, such as lower effective echo spacing and hence mitigated EPI distortions and blurring, and also because of the longer image reconstruction times associated with SMS-EPI data, here data acquisition was performed without the use of SMS in favor of in-plane acceleration using FLEET-ACS and GRAPPA. (Acquisition of subsequent datasets including the Life Span dataset utilized a more newly implemented sequence combining SMS-EPI with FLEET-ACS and GRAPPA, see below.)

In each subject, dMRI data was collected with 4 different b-values (i.e., 4 shells): 1000 s/mm2 (64 directions), 3000 s/mm2 (64 directions), 5000 s/mm2 (128 directions), and 10000 s/mm2 (256 directions). Different b-values were achieved by varying the diffusion gradient amplitudes, while the gradient pulse duration (▯) and diffusion time (▯) were held constant. These b-values were chosen to provide some overlap with the HCP data from WU-Minn consortium, while at the same time push to the limit of diffusion weighting within the constraints of SNR and acquisition time. On determining the number of directions in each shell, in general, data with higher b-value were acquired with more DW directions to capture the increased ratio of high angular frequency components in the MR signal, and to compensate for the SNR loss due to increased diffusion weighting. Also, the number of directions in each shell were selected to meet the typical requirements of popular single shell and multi-shell analysis methods (e.g. q-ball imaging (Descoteaux et al., 2007; Tuch, 2004; Tuch et al., 2002; Tuch et al., 2003), spherical deconvolution (Anderson, 2005; Dell’Acqua et al., 2007; Tournier et al., 2004), ball and stick model (Behrens et al., 2007; Jbabdi et al., 2012), multi-shell q-ball imaging (Aganj et al., 2010; Yeh et al., 2010), diffusion propagator imaging (Descoteaux et al., 2009, 2011), etc.), and to keep the total acquisition time feasible in healthy control subjects.

The diffusion sensitizing direction sets were specifically designed so that the 64-direction set is a subset of the 128-direction set, which is again a subset of the 256-direction set. The initial 64 directions were calculated with the electro-static repulsion method (Caruyer et al., 2013; Jones et al., 1999). With these 64 directions fixed, another 64 directions were added as unknowns, and an optimized 128-direction set was calculated by adjusting the added 64 directions using electrostatic repulsion optimization. With these 128 directions fixed, the 256-direction set was generated using the same method. As such, all 4 shells share the same 64 directions; the b = 5000 s/mm2 shell and the b= 10000 s/mm2 shell share the same 128 directions.

During data acquisition, the diffusion sensitizing directions with approximately opposite polarities were played in pairs to counter-balance the eddy current effects induced by switching the diffusion weighting gradient on and off. Each run started with acquiring a non-DW image (b=0), and one non-DW image was collected every 13 DW images thereafter. Therefore 552 image volumes were collected in total, including 512 DW and 40 non-DW volumes for each subject.

### 2.3 DTI Data Pre-processing and Quality Control

The MGH-USC HCP team completed their basic imaging data preprocessing, with software tools in Free surfer and FSL, which includes (i) Gradient nonlinearity correction, (ii) Motion correction, (iii) Eddy current correction, (iv) b-vectors.

All DTI data were corrected for gradient nonlinearity distortions offline (Glasser et al., 2013; Jovicich et al., 2006). Diffusion data was further corrected for head motion and eddy current artifacts. Specifically, the b=0 images interspersed throughout the diffusion scans were used to estimate bulk head movements with respect to the initial time point (the first b=0 image), where rigid transformation was calculated using the boundary-based registration tool in the Free Surfer package V5.3.0 (Greve and Fischl, 2009). For each b=0 image, this transformation was then applied to itself and the following 13 diffusion weighted images to correct for head motion. Data of all 4 b-values were concatenated (552 image volumes in total) and passed into the EDDY tool (Andersson et al., 2012) for eddy current distortion correction and residual head motion estimates. The rigid rotational components of the motion estimates were then used to adjust the diffusion gradient table for later data reconstruction purposes.

Both unprocessed and minimally pre-processed data are available for download in compressed NIfTI format. The measured gradient field nonlinearity coefficients are protected by Siemens as proprietary information. Because the gradient nonlinearity correction cannot be performed without this information, the unprocessed data provided have already been corrected for gradient nonlinearity distortions with no other pre-processing performed.

All the anatomical scans (T1w and T2w) were free from gross brain abnormalities as determined by an MGH trained physician. All MRI data (T1w, T2w and DW) went through the process of quality assessment by an MGH faculty member, MGH postdoctoral researcher and an MGH research assistant, all trained in neuroimaging in MGH. More specifically, each dataset was assessed by two raters, who viewed both the unprocessed and minimally pre-processed data volume by volume, and rated in terms of head movements, facial and ear mask coverage (to make sure that brain tissue was not masked off), and eddy current correction results. Finally, a comprehensive grade was given to determine whether a dataset had passed quality control.

### 2.4 SUMMARY OF DATASETS

This data set was originally from the MGH-USC HCP team has released diffusion imaging and structural imaging data acquired from 32 young adults using the customized MGH Siemens 3T Connectome scanner, which has 300 mT/m maximum gradient strength for diffusion imaging. A multishell diffusion scheme was used, and the b-values were 1000, 3000, 5000 and 9950 s/mm2. The number of diffusion sampling directions were 64, 64, 128, and 256. The in-plane resolution was 1.5 mm. The slice thickness was 1.5 mm.

### 2.5 FIBER TRACTOGRAPHY

Fiber tractography is a very elegant method, which can used to delineate individual fiber tracts from diffusion images. The main process of study, uses the “DSI-Studio” software tools for i. Complete preprocessing, ii. Fiber tracking and iii. Analysis.

The imaging data processing helps to convert the pre-processed raw data to .src file format, which will be suitable for the further reconstruction process. The reconstruction of .src files achieved through a software tool. It converts the .src imaging data to .fib file format. Only the .fib files are compatible for fiber tracking. To delineate individual fiber tracts from reconstructed diffusion images (.fib file), using a DSI studio software tools.

**Fig 3.**
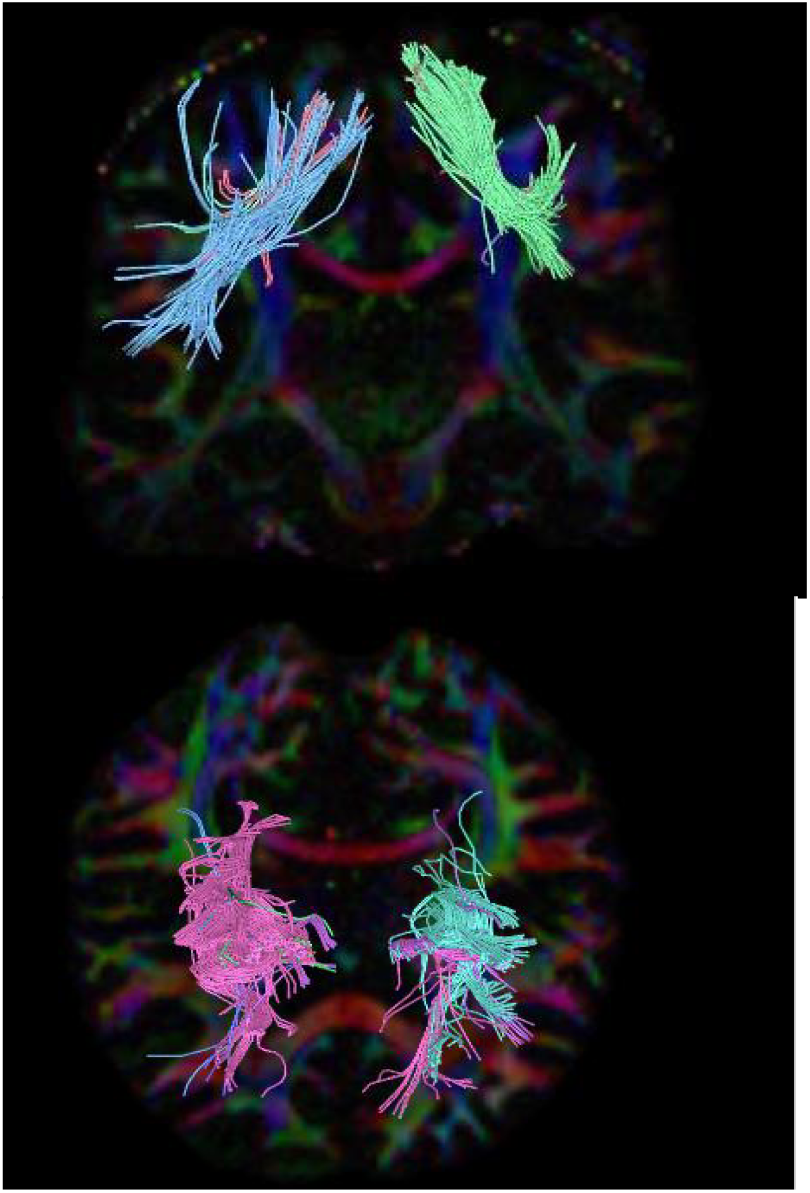
Coronal Section Male Bi – hemispheric Axial Section Female Bi – hemispheric

## RESULTS

### A) Number of tracts

The purpose of our study is deemed to find the neural structures extending between two cortices, connecting them from one end to another end. Fiber tracking uses diffusion tensor to track the fibers along the whole length starting from the seed to end region. The resolution of finding number of tracts helps us to correlate the actual function of dorsal stream visual pathway;

a. Male Right and Left Side
b. Female Right and Left side
c. Right side Male and Female
d. Left side Male and Female

**a) Number of Tracts in Male Subjects Right and Left**

We selected sixteen healthy adults’ male participants for our study having a mean age of 30.4 years.

***Right Side:***

The number of tracts were analyzed in 16 male subjects by tracing the neural structural connectivity between the primary visual cortex to Inferior Temporal Lobe, we found that the male subject with the mean age of 32.5 years has highest number of tracts.

***Left Side:***

The number of tracts were analyzed in 16 male subjects by tracing the neural structural connectivity between the primary visual cortex to Inferior Temporal Lobe, we found that the male subject (1007) with the mean age of 52 has the least number of tracts, (Table 1).

**TABLE 1.** NUMBER OF TRACTS IN MALE RIGHT AND LEFT*

| Subject | SIDE | Number of tracts<br>Male | SIDE | Number of tracts<br>Male |
| --- | --- | --- | --- | --- |
| 1006 | R | 4 | L | 2 |
| 1007 | R | 8 | L | 194 |
| 1008 | R | 8 | L | 4 |
| 1009 | R | 45 | L | 20 |
| 1011 | R | 2 | L | 8 |
| 1013 | R | 25 | L | 11 |
| 1014 | R | 42 | L | 53 |
| 1015 | R | 8 | L | 20 |
| 1016 | R | 10 | L | 38 |
| 1022 | R | 6 | L | 11 |
| 1025 | R | 8 | L | 1 |
| 1027 | R | 16 | L | 18 |
| 1028 | R | 111 | L | 19 |
| 1029 | R | 7 | L | 29 |
| 1030 | R | 2 | L | 31 |
| 1031 | R | 26 | L | 11 |

The $p$ -value is .513463 \*The results were statistically **not** significant $p < .10$

**Fig 1.**
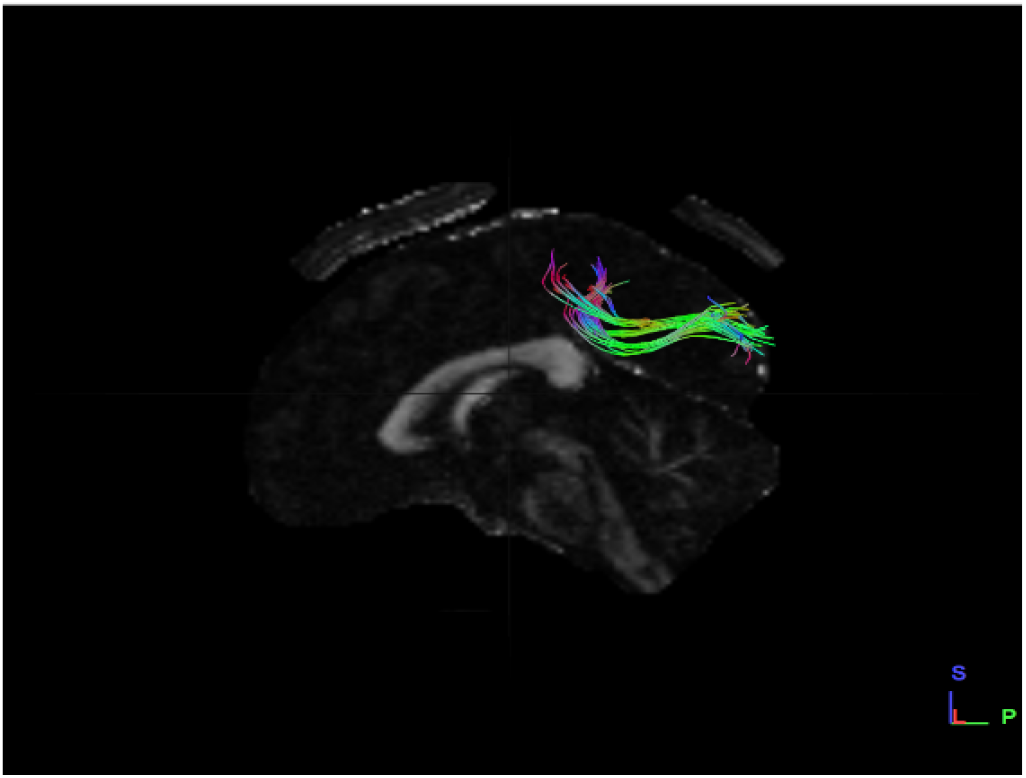
Sagittal Section Male Left Side Dataset

**Fig 2.**
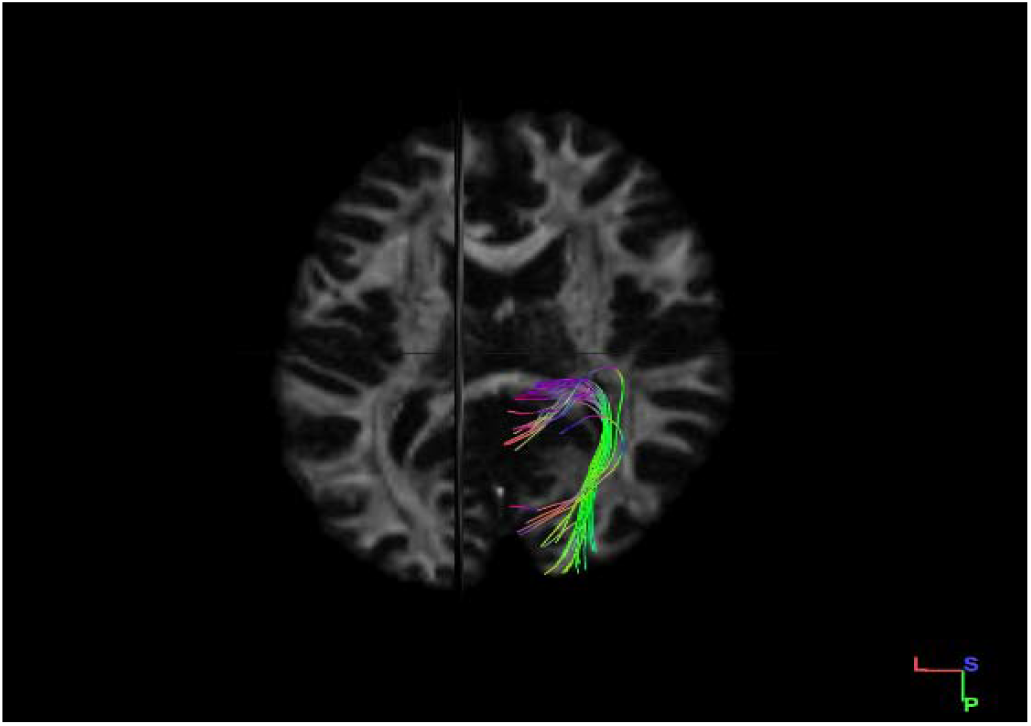
Axial Section Male Right Side Dataset

**<u>Graphical Representation</u>**

**Grap-1.**
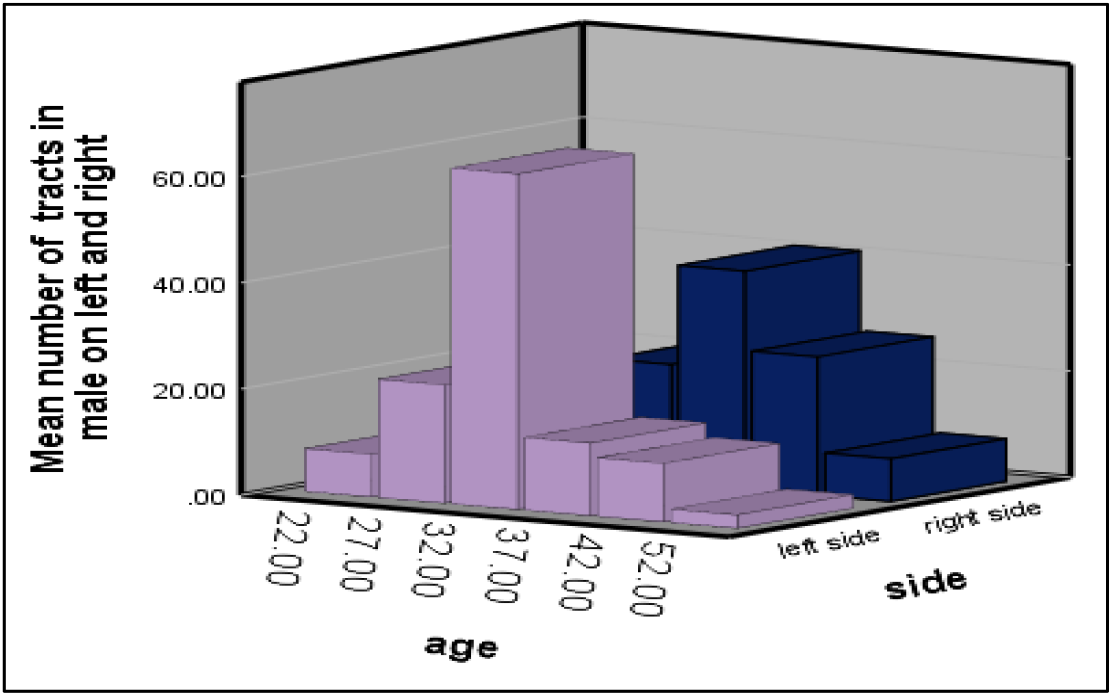
Number of tracts in male subjects on both left and right sides.

We found that on average, the left side has a greater number of tracts than on right side in male subjects.

**b) Number of Tracts in Female Subjects Right and Left**

We selected sixteen healthy adult female participants for our study, having a mean age of 30.4 years.

***Right Side:***

The number of tracts were analyzed in 16 female subjects by tracing the neural structural connectivity between the primary visual cortex to Inferior Temporal Lobe, we found the male subject (1033) with the mean age of 42 years having highest number of tracts.

***Left Side:***

The number of tracts were analyzed in 16 female subjects by tracing the neural structural connectivity between the primary visual cortex to Inferior Temporal Lobe, we found the female subject (1001) with the mean age of 22 having least number of tracts, (Table 2).

**TABLE 2.** NUMBER OF TRACTS IN FEMALE RIGHT AND LEFT*

| Subject | SIDE | Number of tracts | SIDE | Number of tracts |
| --- | --- | --- | --- | --- |
| 1001 | R | 17 | L | 1 |
| 1002 | R | 69 | L | 50 |
| 1003 | R | 66 | L | 316 |
| 1004 | R | 4 | L | 35 |
| 1005 | R | 55 | L | 14 |
| 1010 | R | 70 | L | 6 |
| 1012 | R | 17 | L | 2 |
| 1017 | R | 16 | L | 67 |
| 1019 | R | 4 | L | 39 |
| 1021 | R | 13 | L | 32 |
| 1023 | R | 4 | L | 79 |
| 1024 | R | 7 | L | 6 |
| 1026 | R | 7 | L | 62 |
| 1032 | R | 14 | L | 14 |
| 1033 | R | 405 | L | 10 |
| 1035 | R | 9 | L | 9 |

**Fig 3.**
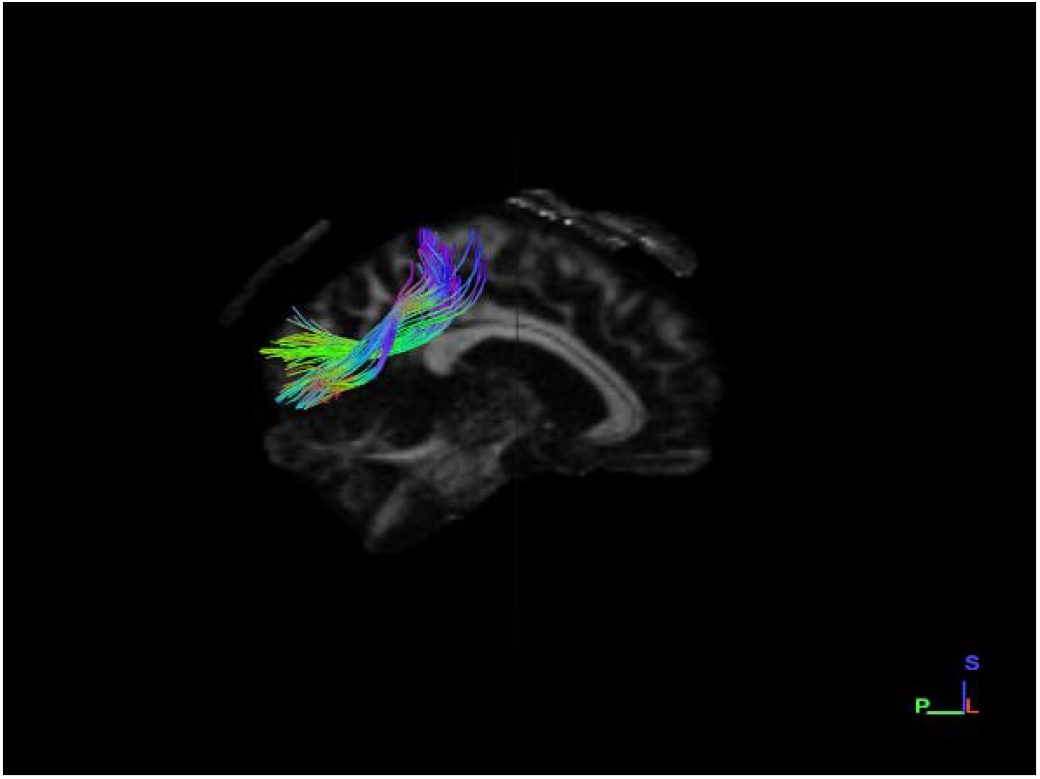
Sagittal Section Female Right Side Dataset

**Fig 4.**
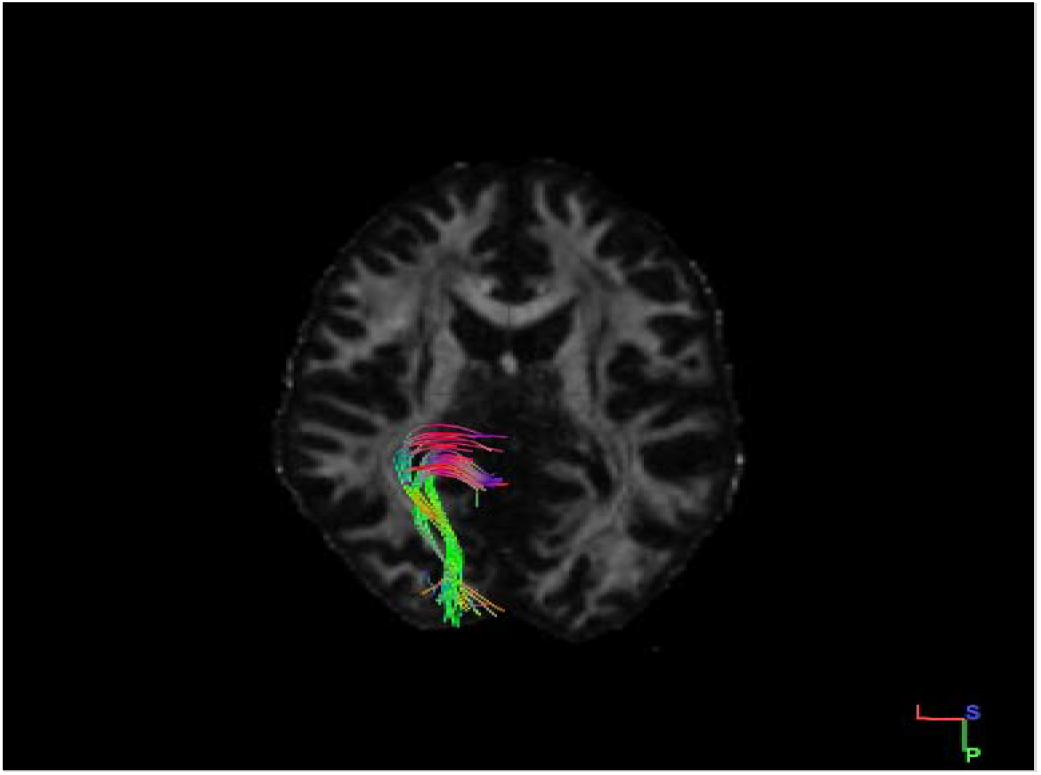
Axial Section Female Left Side Dataset

**<u>Graphical Representation</u>**

**Grap-2.**
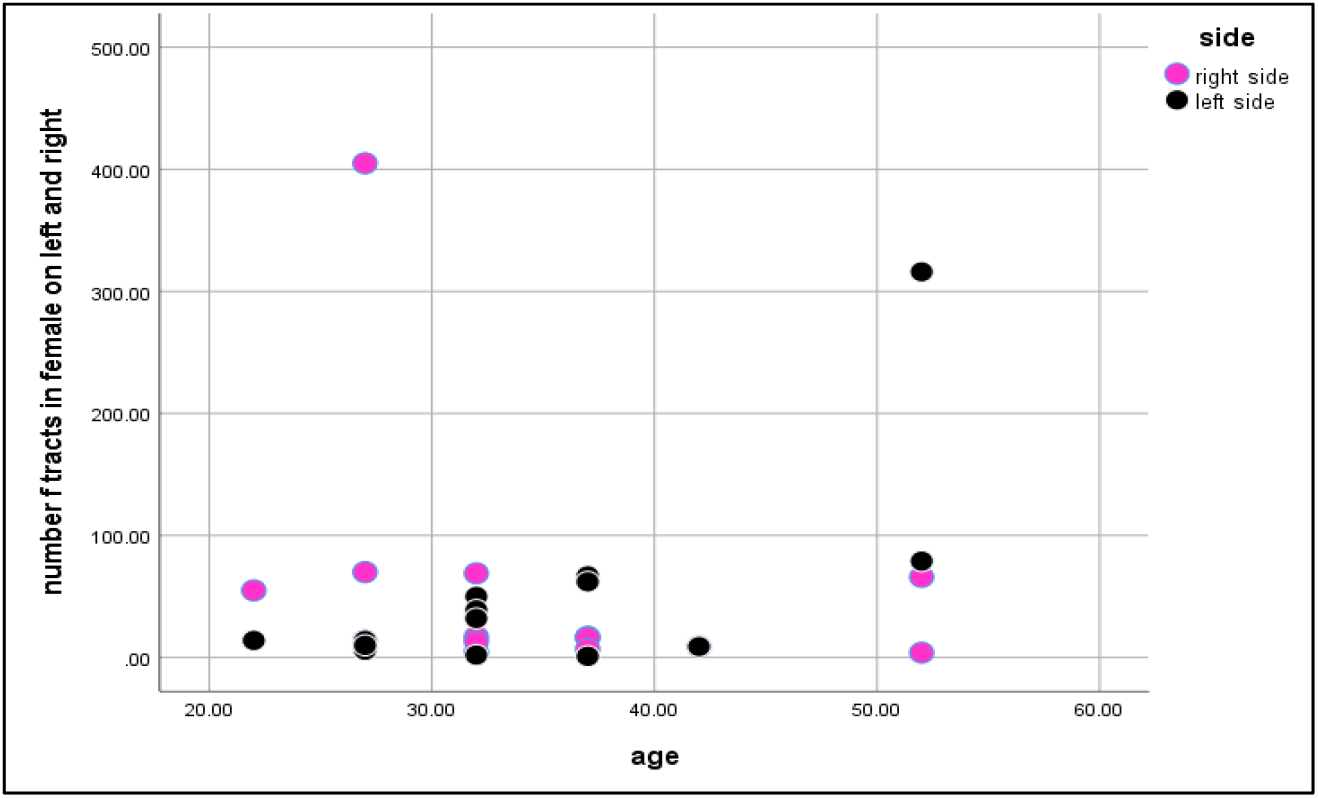
Number of tracts in female subjects on both left and right sides.

We found the right side having greater number of tracts than left side in female subjects.

**TABLE 3.** NUMBER OF TRACTS IN MALE AND FEMALE RIGHT*

| Female Datasets | Number of tracts Right | Male Datasets | Number of tracts Right |
| --- | --- | --- | --- |
| 1001 | 17 | 1006 | 4 |
| 1002 | 69 | 1007 | 8 |
| 1003 | 66 | 1008 | 8 |
| 1004 | 4 | 1009 | 45 |
| 1005 | 55 | 1011 | 2 |
| 1010 | 70 | 1013 | 25 |
| 1012 | 17 | 1014 | 42 |
| 1017 | 16 | 1015 | 8 |
| 1019 | 4 | 1016 | 10 |
| 1021 | 13 | 1022 | 6 |
| 1023 | 4 | 1025 | 8 |
| 1024 | 7 | 1027 | 16 |
| 1026 | 7 | 1028 | 111 |
| 1032 | 14 | 1029 | 7 |
| 1033 | 405 | 1030 | 2 |
| 1035 | 9 | 1031 | 26 |

The $p$ -value is .280034. The result was statistically *not* significant at $p < .10$

**Fig 4.**
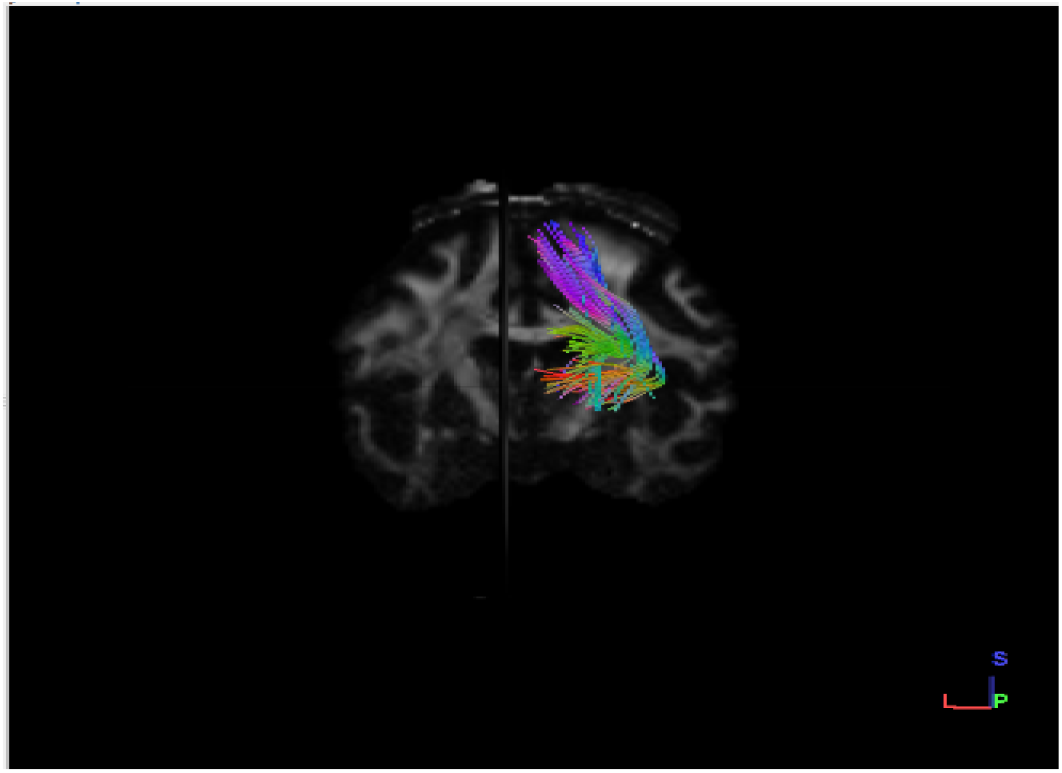
Coronal section female right side dataset

**Figure 5.**
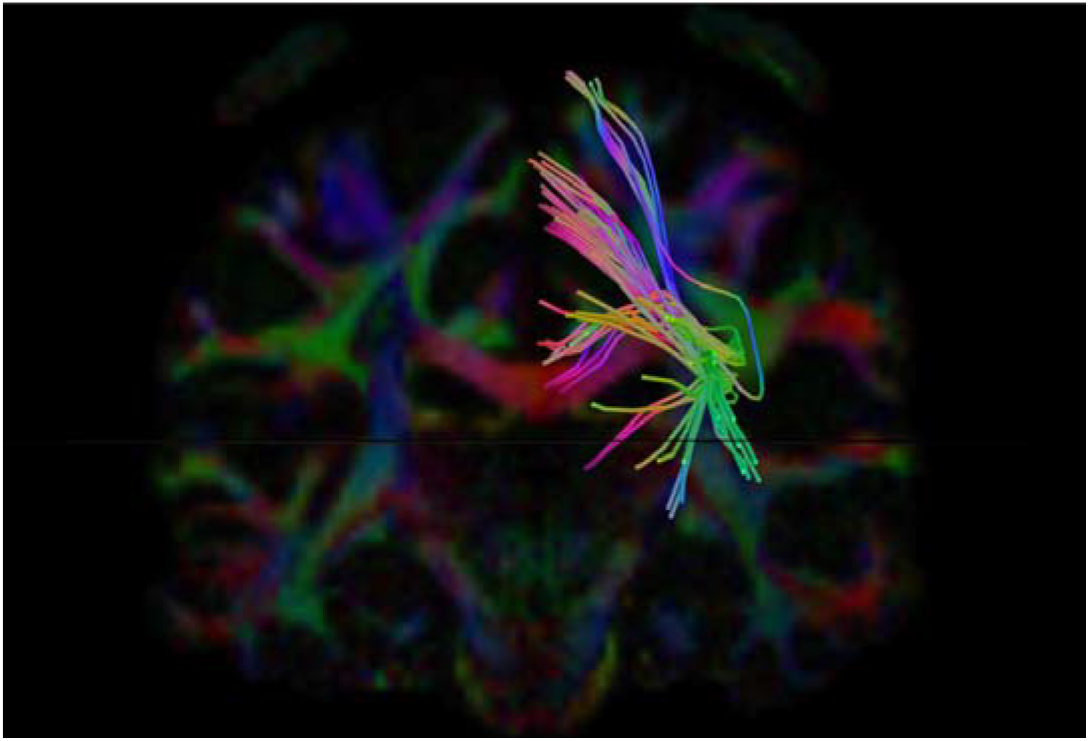
Coronal section male right side dataset

We found that female subjects have greater number of fibers on the right side than in male subjects.

**TABLE 4.**
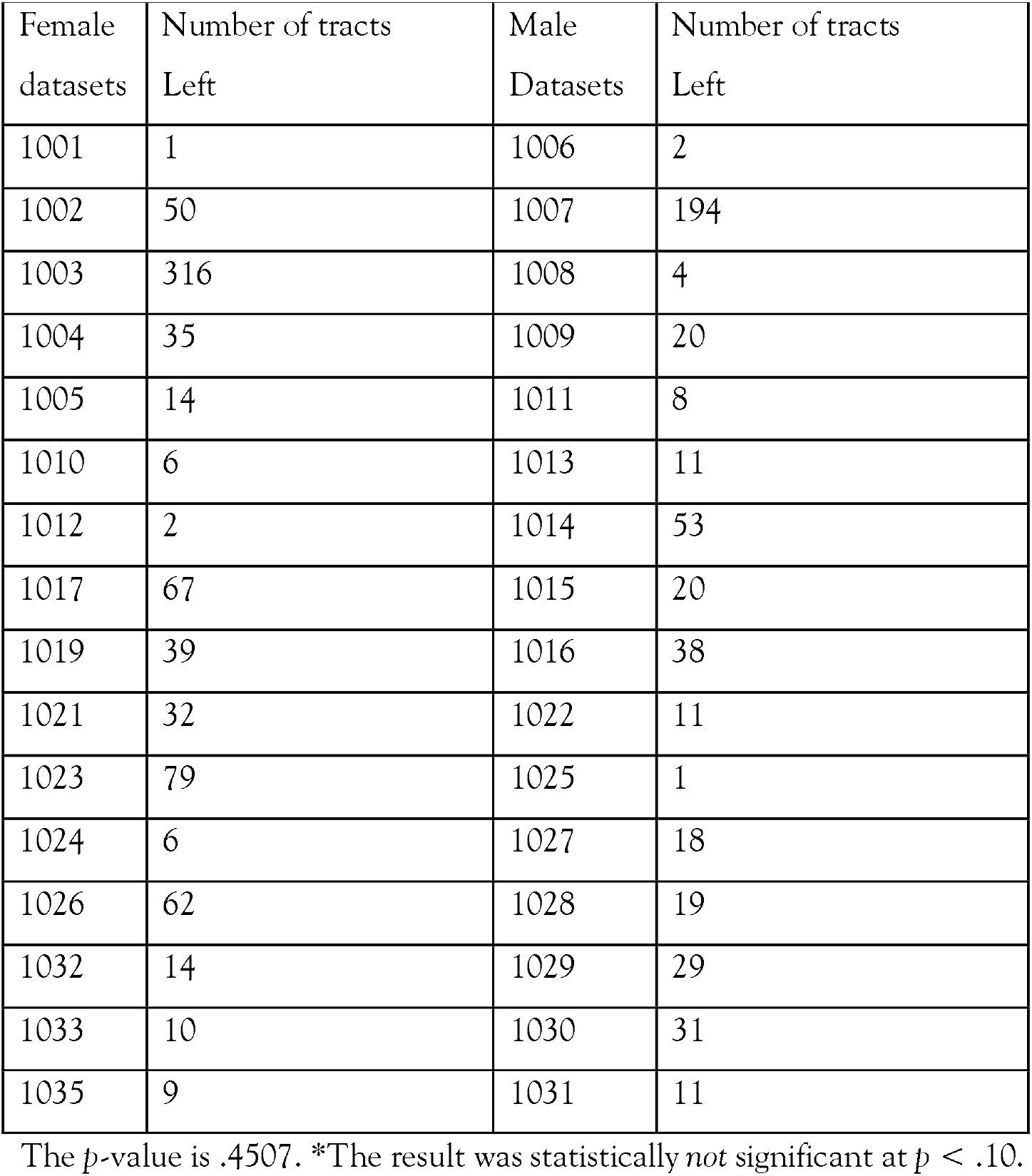
NUMBER OF TRACTS IN MALE AND FEMALE LEFT*

| Female datasets | Number of tracts Left | Male Datasets | Number of tracts Left |
| --- | --- | --- | --- |
| 1001 | 1 | 1006 | 2 |
| 1002 | 50 | 1007 | 194 |
| 1003 | 316 | 1008 | 4 |
| 1004 | 35 | 1009 | 20 |
| 1005 | 14 | 1011 | 8 |
| 1010 | 6 | 1013 | 11 |
| 1012 | 2 | 1014 | 53 |
| 1017 | 67 | 1015 | 20 |
| 1019 | 39 | 1016 | 38 |
| 1021 | 32 | 1022 | 11 |
| 1023 | 79 | 1025 | 1 |
| 1024 | 6 | 1027 | 18 |
| 1026 | 62 | 1028 | 19 |
| 1032 | 14 | 1029 | 29 |
| 1033 | 10 | 1030 | 31 |
| 1035 | 9 | 1031 | 11 |

The $p$ -value is .4507. \*The result was statistically *not* significant at $p < .10$ .

**Fig 6.**
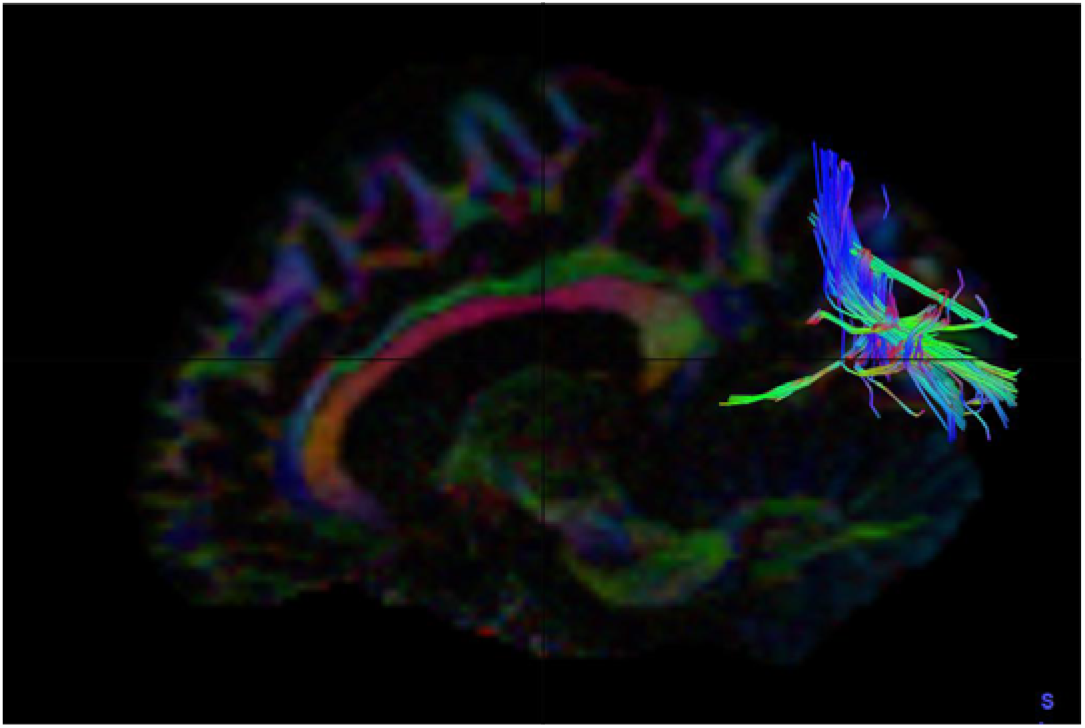
Sagittal section female left side dataset

**Fig 7.**
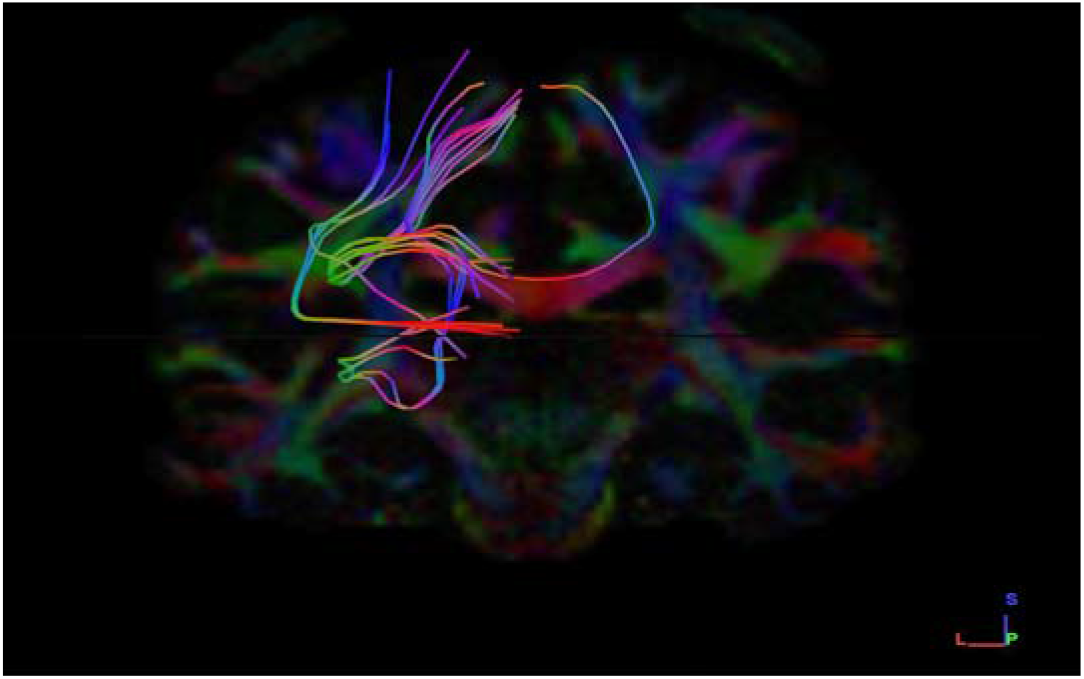
Coronal section male left side dataset

**<u>Graphical Representation</u>**

**Grap-3.**
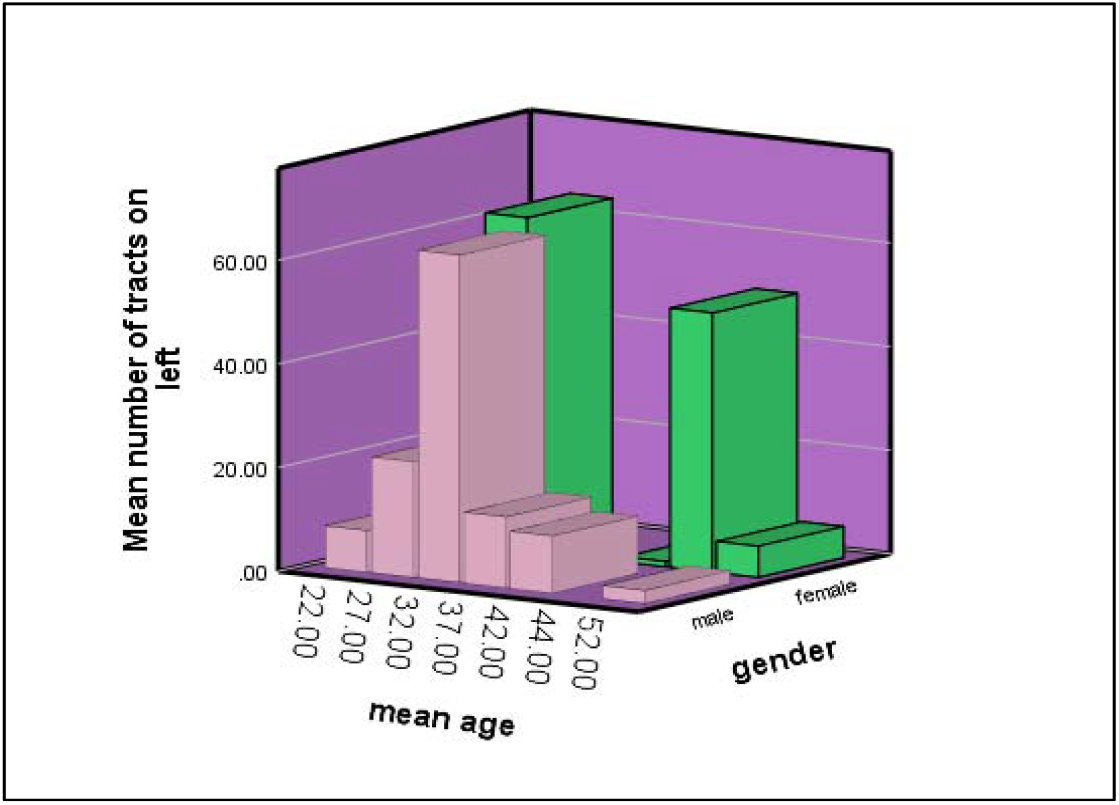
Number of tracts in both male and female subjects left side.

We found that female subjects have greater number of fibers on left side than in male subjects.

### B) Volume of tracts

Tract volume is the total volume occupied by fibres collectively. Total volume is otherwise called the total number of voxels that were assigned as containing neural fibers. The comparison of volume of Fibers is mostly done between the seed and the end region under four parameters;

a. Male Right and Left Side
b. Female Right and Left side
c. Right side Male and Female
d. Left side Male and Female

**a) Volume of Tracts in Males, both Right and Left Side**

We selected sixteen healthy adult male participants for our study having a mean age of 30.4 years.

***Right Side:***

The Volume of tracts were analyzed in 16 male subjects by tracing the neural structural connectivity between the primary visual cortex to Inferior Temporal Lobe, we found that the male subject (1028) with the mean age of 32.5 years having highest volume of tracts.

***Left Side:***

The number of tracts were analyzed in 16 male subjects by tracing the neural structural connectivity between the primary visual cortex to Inferior Temporal Lobe, we found that the male subject (1025) with the mean age of 37 having least volume of tracts, (Table 5).

**TABLE 5.**
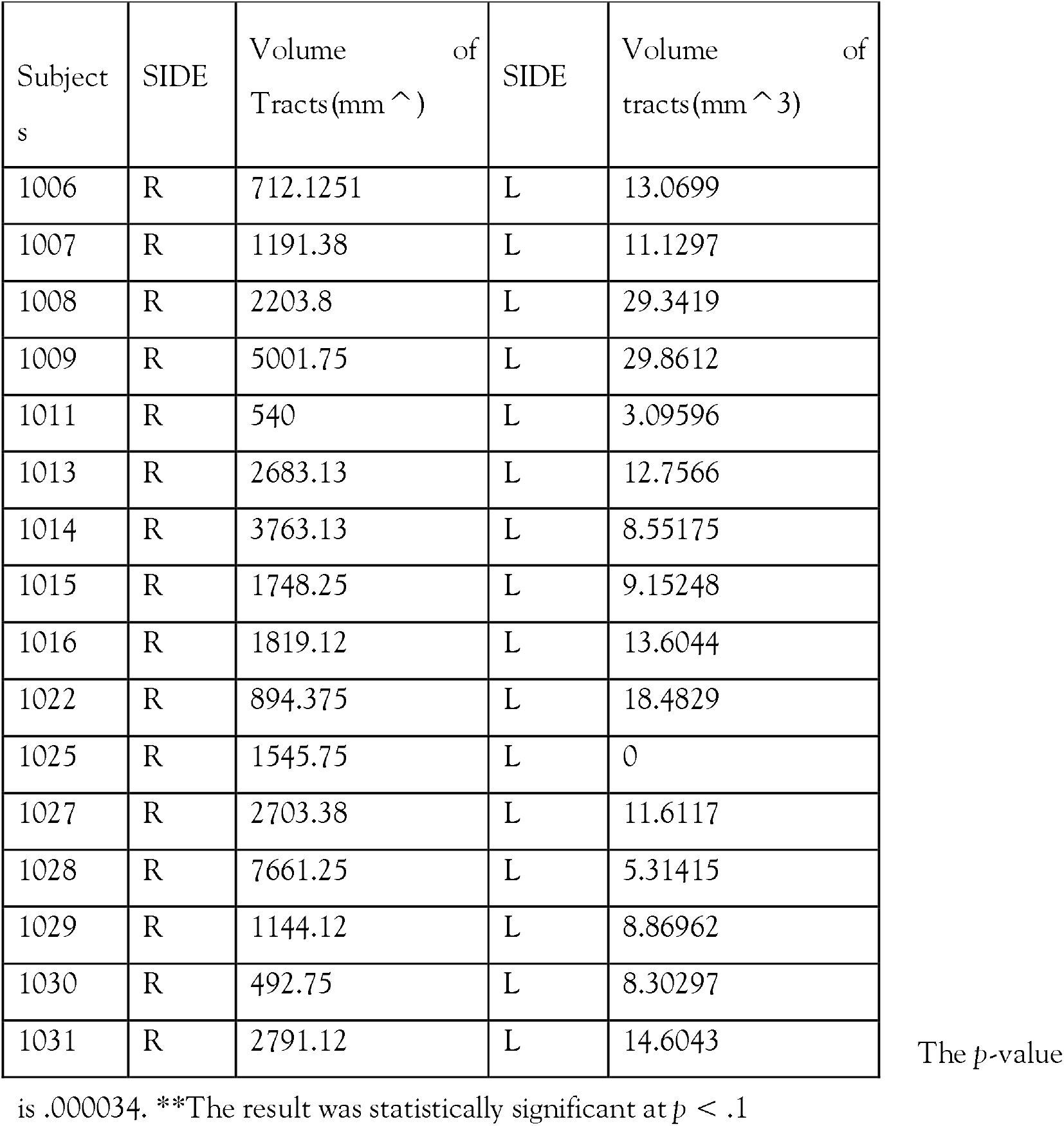
VOLUME OF TRACTS IN MALE RIGHT AND LEFT**

| Subject<br>s | SIDE | Volume<br>of<br>Tracts(mm ^ ) | SIDE | Volume<br>of<br>tracts(mm ^ 3) |
| --- | --- | --- | --- | --- |
| 1006 | R | 712.1251 | L | 13.0699 |
| 1007 | R | 1191.38 | L | 11.1297 |
| 1008 | R | 2203.8 | L | 29.3419 |
| 1009 | R | 5001.75 | L | 29.8612 |
| 1011 | R | 540 | L | 3.09596 |
| 1013 | R | 2683.13 | L | 12.7566 |
| 1014 | R | 3763.13 | L | 8.55175 |
| 1015 | R | 1748.25 | L | 9.15248 |
| 1016 | R | 1819.12 | L | 13.6044 |
| 1022 | R | 894.375 | L | 18.4829 |
| 1025 | R | 1545.75 | L | 0 |
| 1027 | R | 2703.38 | L | 11.6117 |
| 1028 | R | 7661.25 | L | 5.31415 |
| 1029 | R | 1144.12 | L | 8.86962 |
| 1030 | R | 492.75 | L | 8.30297 |
| 1031 | R | 2791.12 | L | 14.6043 |

The *p*-value
is .000034. \*\*The result was statistically significant at $p < .1$

**Fig 8.**
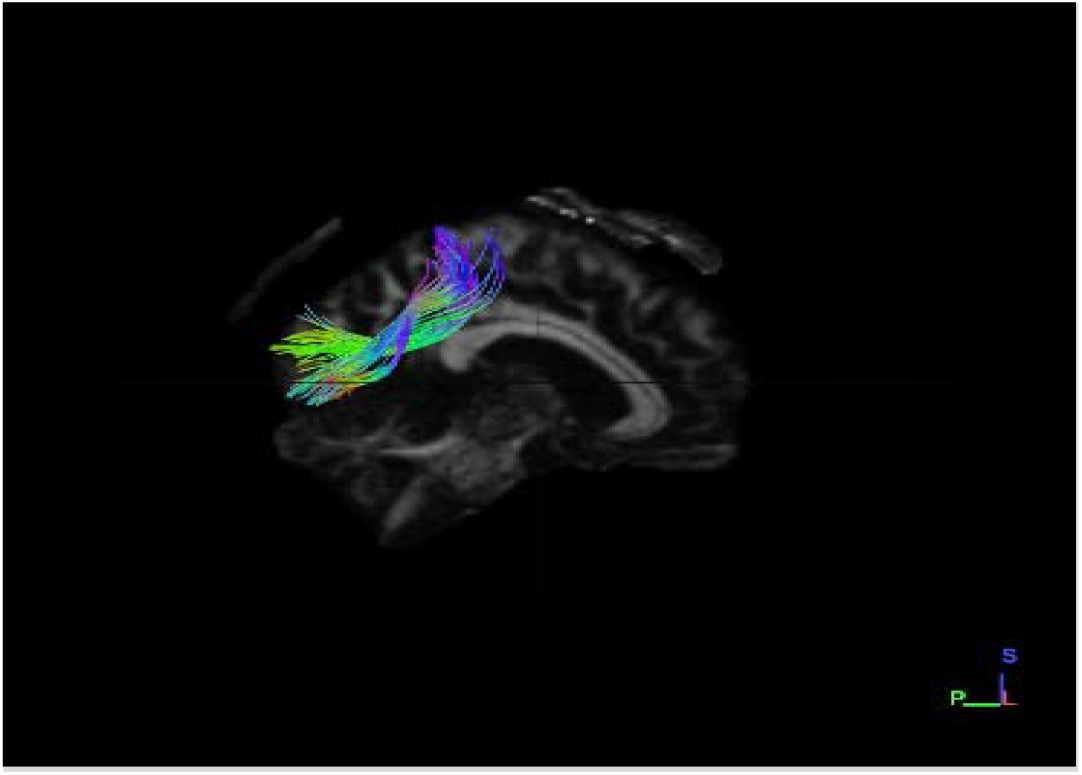
Sagittal section male right side dataset

**Fig 9.**
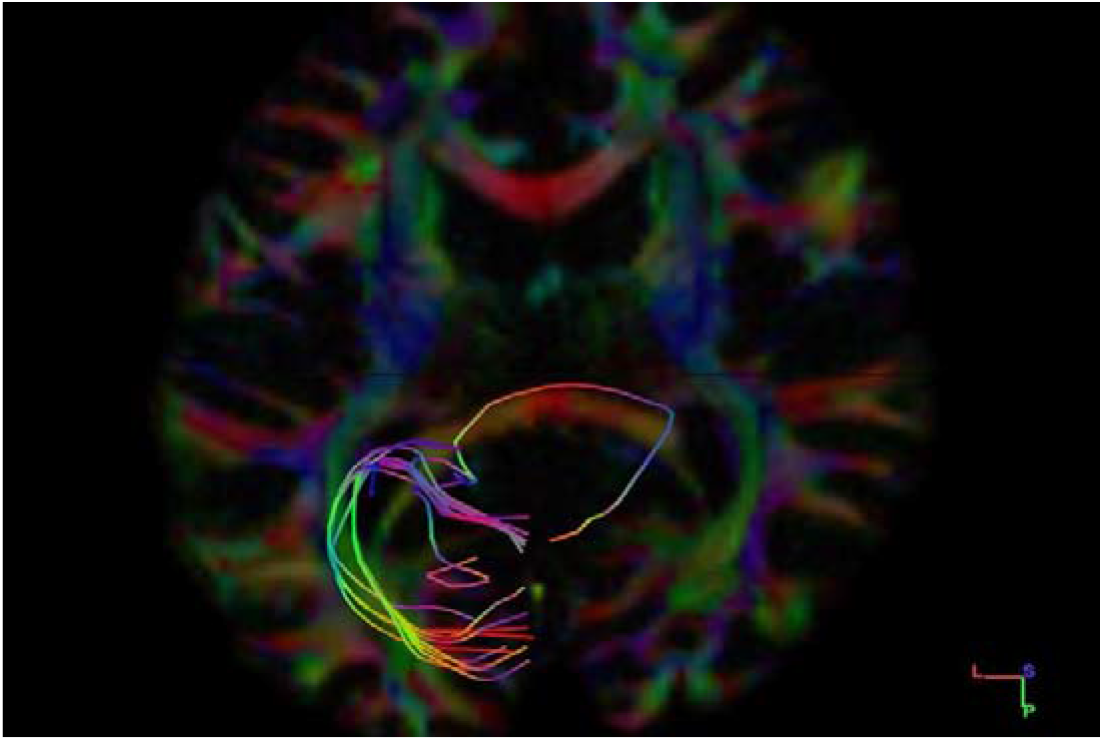
Axial section male left side dataset

**<u>Graphical Representation</u>**

**Grap-4.**
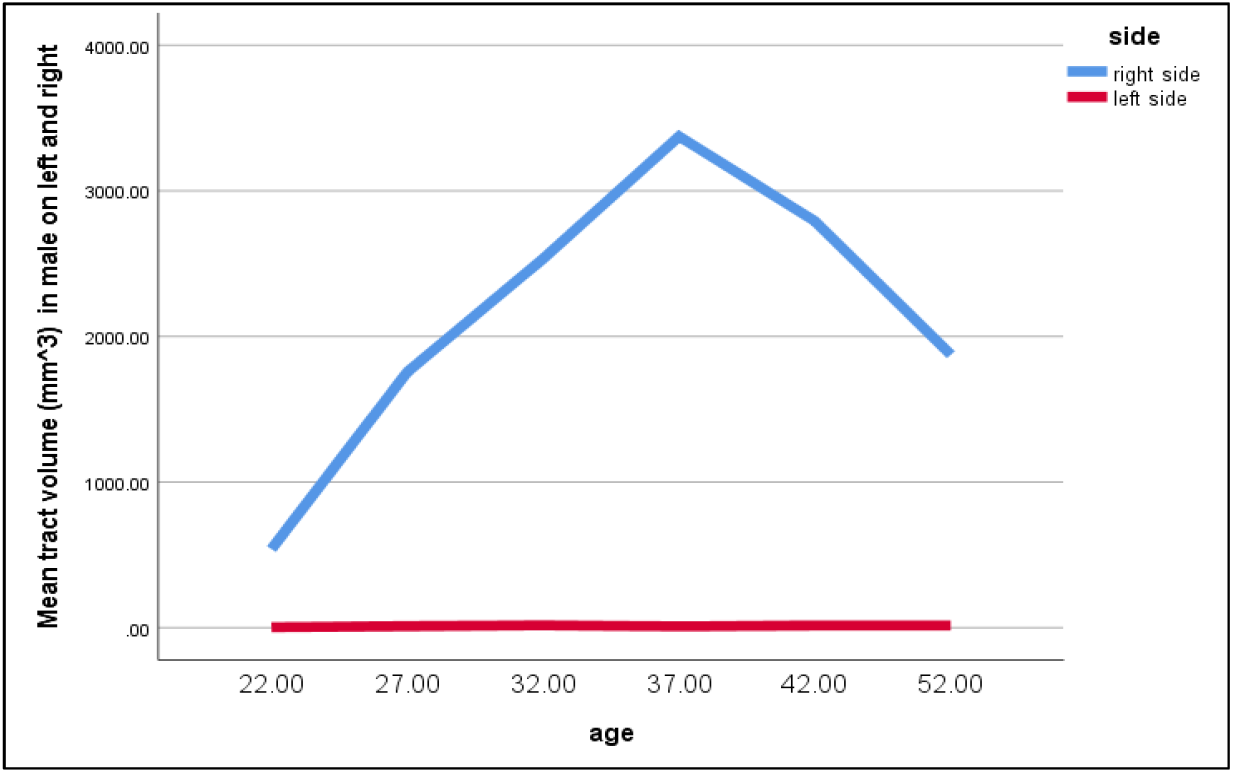
Tract volume in male subjects on left and right sides.

We found that on average, the right side is having greater volume of tracts than on left side in male subjects.

**b) Volume of Tracts in Females on both Right and Left Side**

We selected sixteen healthy adult female participants for our study, having a mean age of 30.4 years.

***Right Side:***

The Volume of tracts were analyzed in 16 female subjects by tracing the neural structural connectivity between the primary visual cortex to Inferior Temporal Lobe, we found that the female subject (1010) with the mean age of 27 years having highest number of tracts.

***Left Side:***

The volume of tracts was analyzed in 16 female subjects by tracing the neural structural connectivity between the primary visual cortex to Inferior Temporal Lobe, we found that the female subject (1001) with the mean age of 41.4 having least volume of tracts, (Table-6).

**TABLE 6.** VOLUME OF TRACTS IN FEMALE RIGHT AND LEFT**

| Subjects | SIDE | Volume of Tracts(mm ^ ) | SIDE | Volume of tracts(mm ^ 3) |
| --- | --- | --- | --- | --- |
| 1001 | R | 2072.25 | L | 0 |
| 1002 | R | 4556.25 | L | 59.3764 |
| 1003 | R | 4910.63 | L | 17.5082 |
| 1004 | R | 729 | L | 7.0101 |
| 1005 | R | 5258.25 | L | 5.26428 |
| 1010 | R | 5602.5 | L | 14.3357 |
| 1012 | R | 2254.5 | L | 12.2234 |
| 1017 | R | 2426.62 | L | 9.29758 |
| 1019 | R | 617.625 | L | 7.46943 |
| 1021 | R | 1967.63 | L | 12.2787 |
| 1023 | R | 1029.38 | L | 8.52719 |
| 1024 | R | 1184.63 | L | 42.722 |
| 1026 | R | 1410.75 | L | 11.6172 |
| 1032 | R | 1572.75 | L | 86.3565 |
| 1033 | R | 11454.7 | L | 34.9819 |
| 1035 | R | 1275.75 | L | 87.4314 |

The $p$ -value is .000179. \*\*The result was statistically significant at $p < .10$

**Fige 10.**
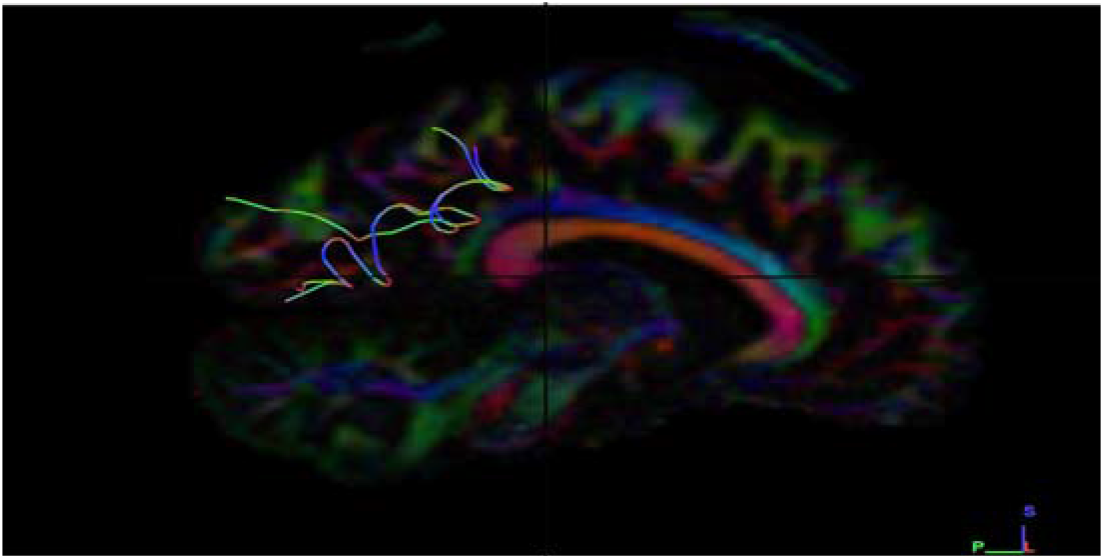
Sagittal section female right side dataset (1010)

**Fige 11.**
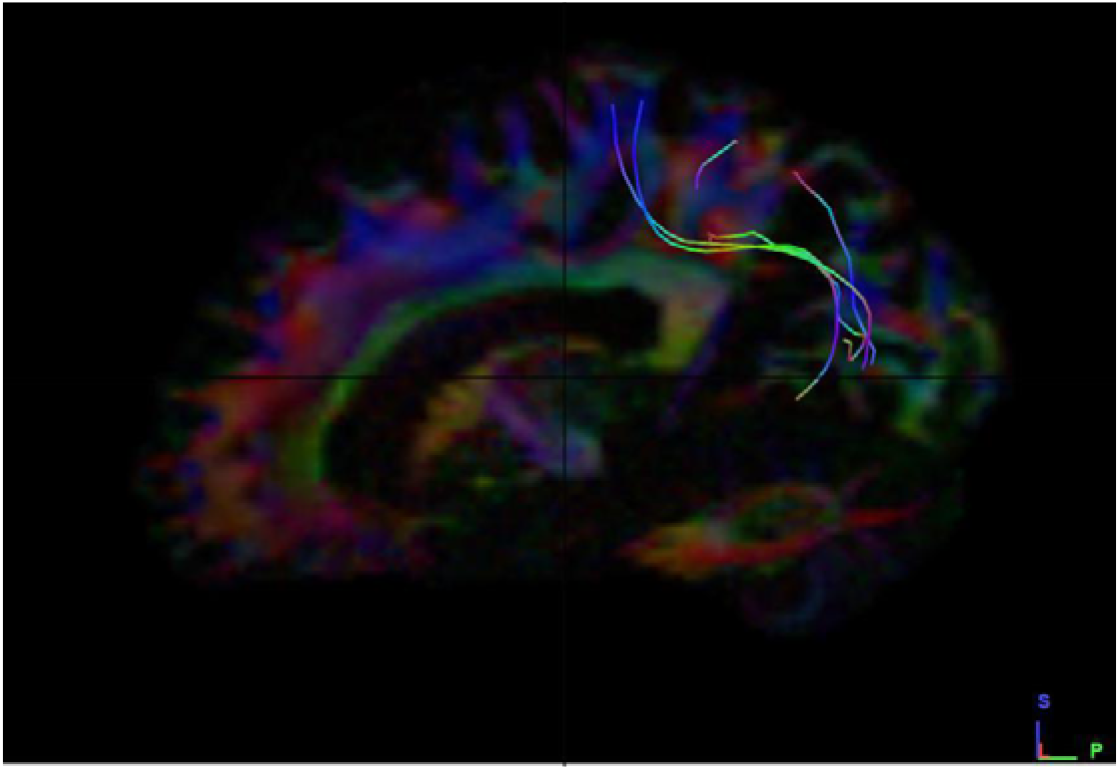
Sagittal section female left side dataset (1010)

**<u>Graphical Representation</u>**

**Graph-5.**
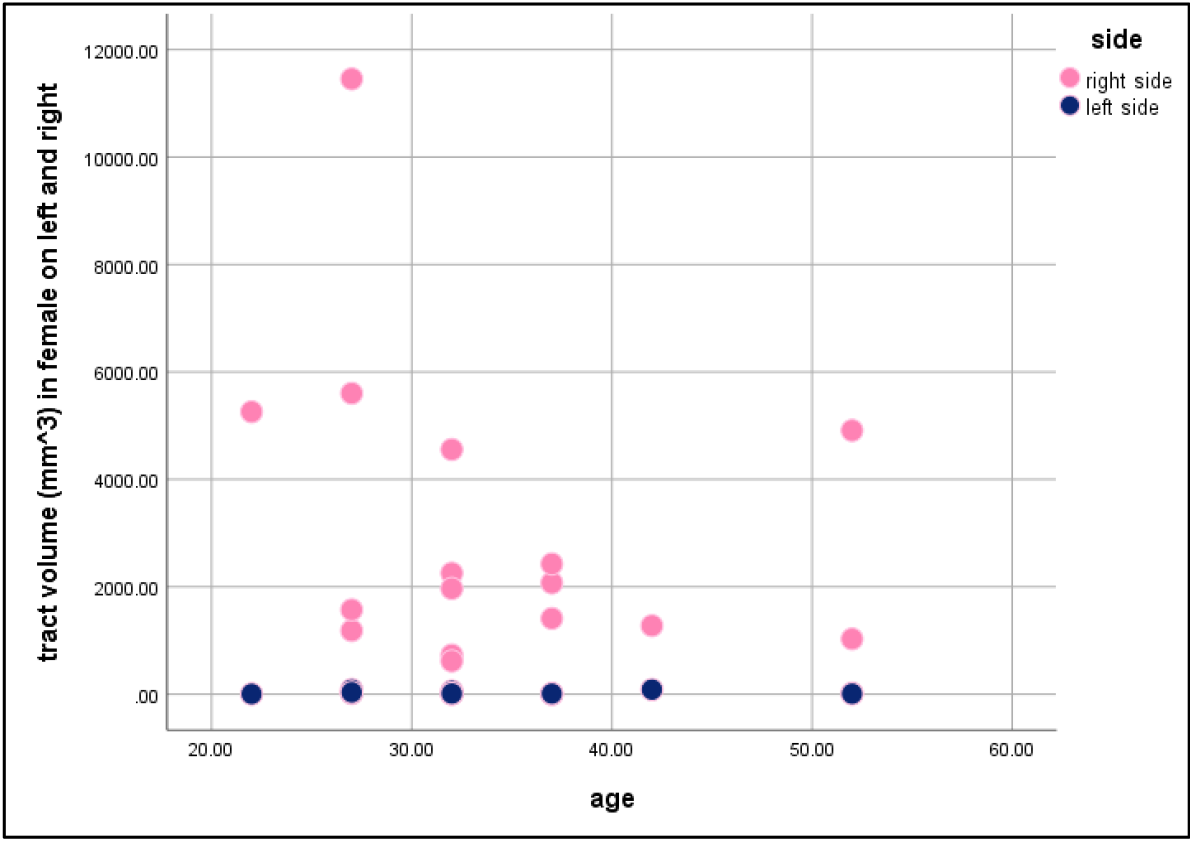
Volume of tracts in female subjects on both left and right sides.

We found that on average, the right side has greater volume of tracts than on left side in female subjects.

**c) Volume of Tracts in both Male and Female Left Side**

We selected sixteen healthy adult male participants for our study, having a mean age of 30.4 years.

***Female:***

The volume of tracts was analyzed in 16 female subjects by tracing the neural structural connectivity between the primary visual cortex to Inferior Temporal Lobe, we found that the female subject (1032) with the mean age of 27 years has highest volume of tracts.

***Male:***

The volume of tracts were analyzed in 16 male subjects by tracing the neural structural connectivity between the primary visual cortex to Inferior Temporal Lobe, we found that the male subject (1023) with the mean age of 27 has least volume of tracts, (Table 7).

**TABLE 7.** VOLUME OF TRACTS IN MALE AND FEMALE LEFT**

| Female<br>Datasets | Volume<br>of<br>Tracts(mm ^ 3) | Male<br>Datasets | Volume of tracts(mm ^ 3) |
| --- | --- | --- | --- |
| 1001 | 0 | 1006 | 13.0699 |
| 1002 | 59.3764 | 1007 | 11.1297 |
| 1003 | 17.5082 | 1008 | 29.3419 |
| 1004 | 7.0101 | 1009 | 29.8612 |
| 1005 | 5.26428 | 1011 | 3.09596 |
| 1010 | 14.3357 | 1013 | 12.7566 |
| 1012 | 12.2234 | 1014 | 8.55175 |
| 1017 | 9.29758 | 1015 | 9.15248 |
| 1019 | 7.46943 | 1016 | 13.6044 |
| 1021 | 12.2787 | 1022 | 18.4829 |
| 1023 | 8.52719 | 1025 | 0 |
| 1024 | 42.722 | 1027 | 11.6117 |
| 1026 | 11.6172 | 1028 | 5.31415 |
| 1032 | 86.3565 | 1029 | 8.86962 |
| 1033 | 34.9819 | 1030 | 8.30297 |
| 1035 | 87.4314 | 1031 | 14.6043 |

The $p$ -value is .073828. \*\*The result was statistically significant at $p < .10$

**<u>Graphical Representation</u>**

**Grap-6.**
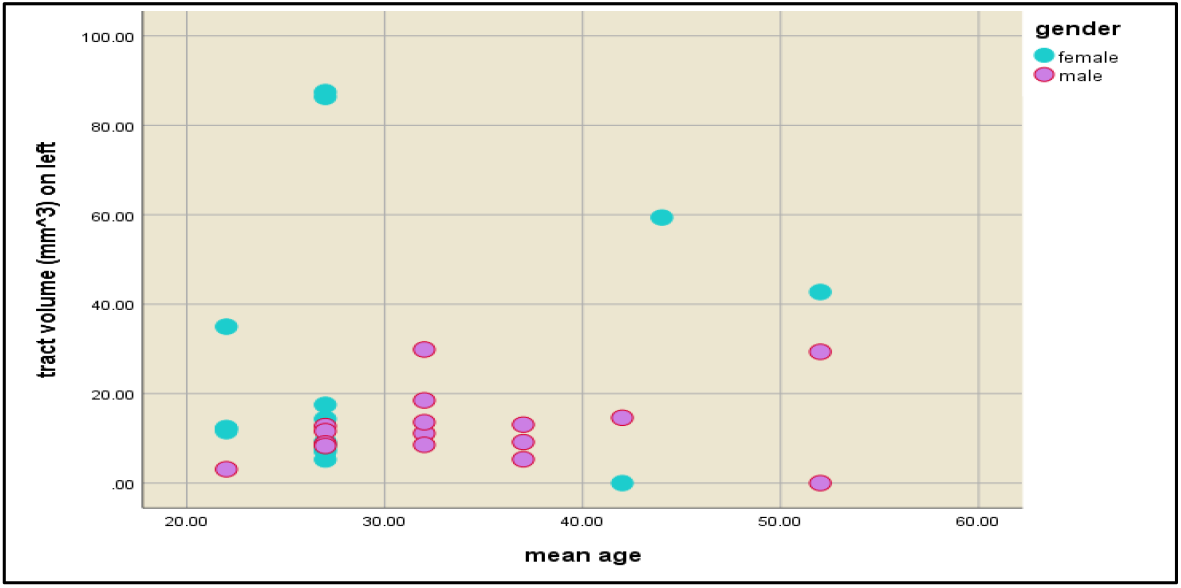
Graphical representation of volume of tracts in male and female subjects on left sides.

We found that female subjects have greater volume of fibres on left side than in male subjects.

**d) Volume of Tracts on both Male and Female Right Side**

We selected sixteen healthy adult male participants for our study having a mean age of 30.4 years.

***Female:***

The volume of tracts was analyzed in 16 female subjects by tracing the neural structural connectivity between the primary visual cortex to Inferior Temporal Lobe, we found that the female subject (1017) with the mean age of 27 years has highest volume of tracts.

***Male:***

The volume of tracts were analyzed in 16 male subjects by tracing the neural structural connectivity between the primary visual cortex to Inferior Temporal Lobe, we found that the male subject (1005) with the mean age of 27 has least volume of tracts, (Table 8).

**TABLE 8.** VOLUME OF TRACTS IN MALE AND FEAMLE RIGHT*

| Female Datasets | Volume of Tracts(mm ^ ) | Male Datasets | Volume of tracts(mm ^ 3) |
| --- | --- | --- | --- |
| 1001 | 2072.25 | 1006 | 712.1251 |
| 1002 | 4556.25 | 1007 | 1191.38 |
| 1003 | 4910.63 | 1008 | 2203.8 |
| 1004 | 729 | 1009 | 5001.75 |
| 1005 | 5258.25 | 1011 | 540 |
| 1010 | 5602.5 | 1013 | 2683.13 |
| 1012 | 2254.5 | 1014 | 3763.13 |
| 1017 | 2426.62 | 1015 | 1748.25 |
| 1019 | 617.625 | 1016 | 1819.12 |
| 1021 | 1967.63 | 1022 | 894.375 |
| 1023 | 1029.38 | 1025 | 1545.75 |
| 1024 | 1184.63 | 1027 | 2703.38 |
| 1026 | 1410.75 | 1028 | 7661.25 |
| 1032 | 1572.75 | 1029 | 1144.12 |
| 1033 | 11454.7 | 1030 | 8.30297 |
| 1035 | 1275.75 | 1031 | 14.6043 |

The $p$ -value is .294117. \*The result was not statistically significant at $p < .10$

**Fig 12.**
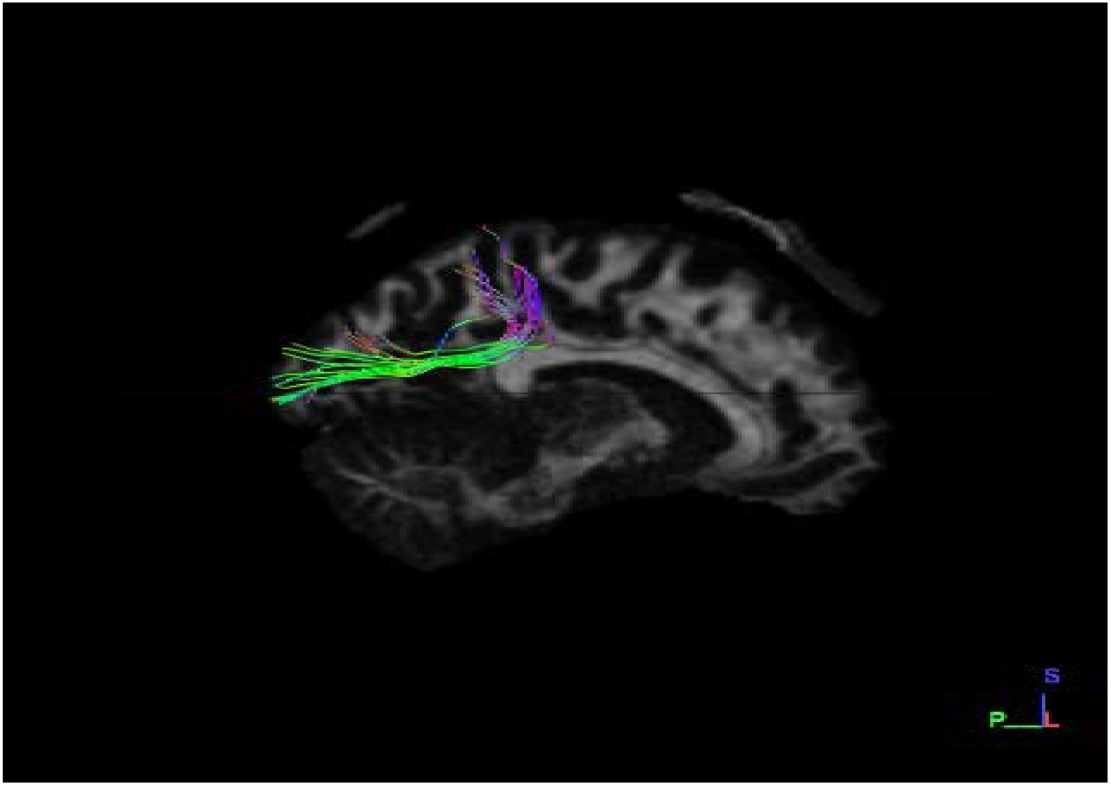
Sagittal section female right side dataset (1017)

**Fig 13.**
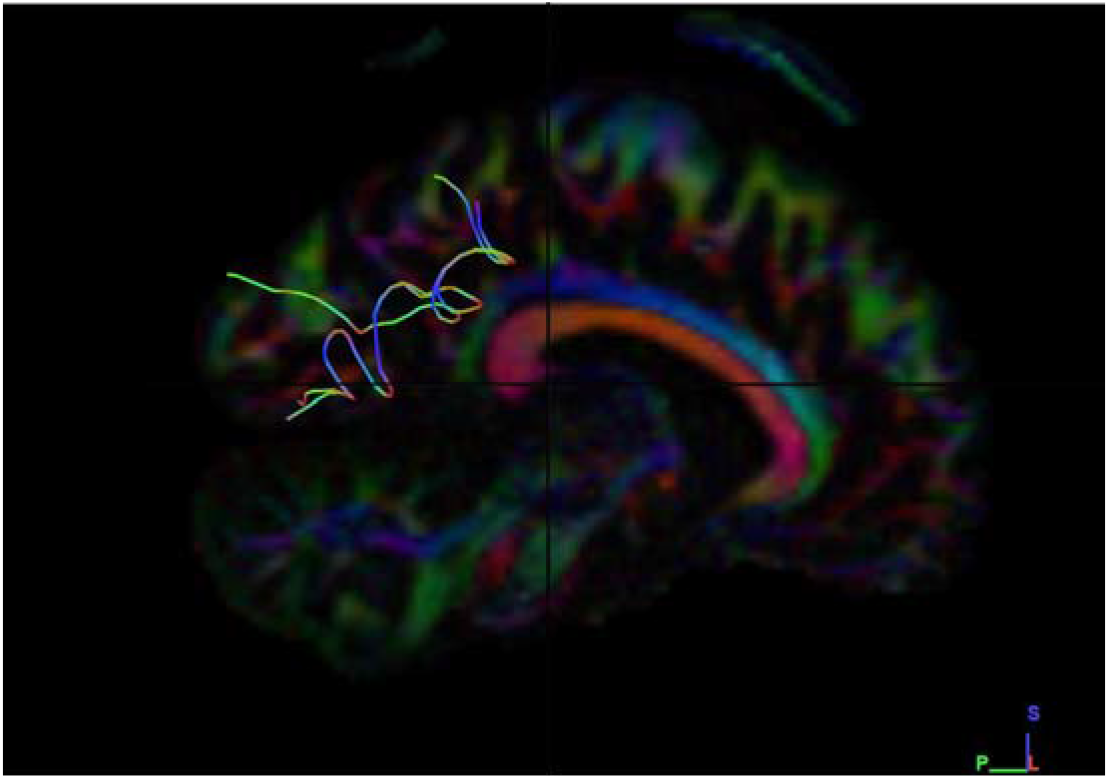
Sagittal section male right side dataset (1017)

**<u>Graphical Representation</u>**

**Graph-7.**
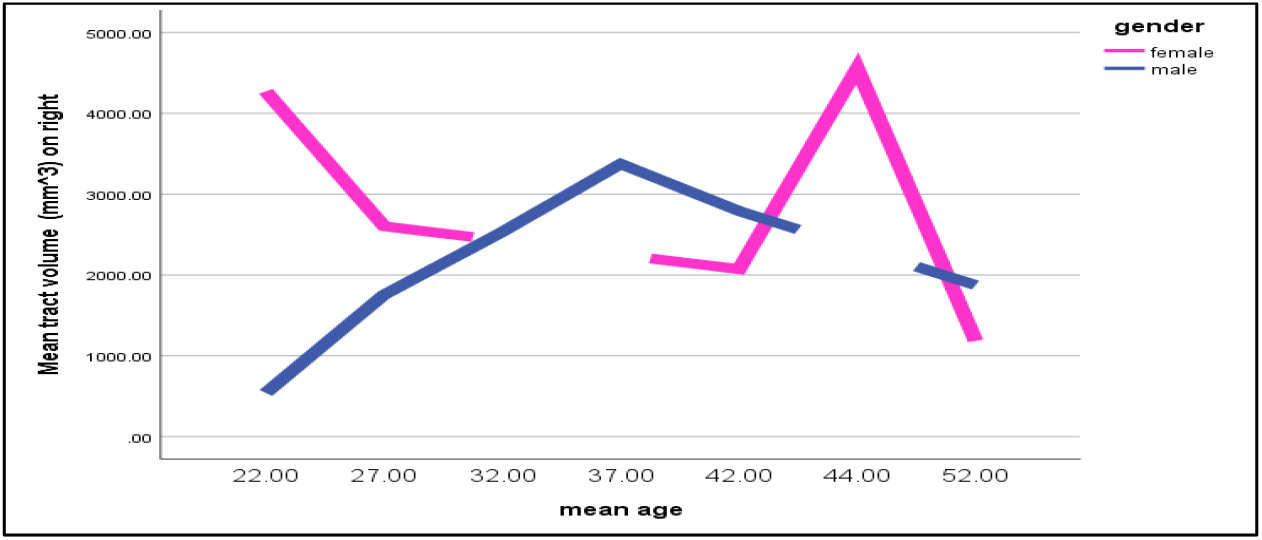
Graphical representation of volume of tracts in male and female subjects on the right side

We found that female subjects have greater volume fibers on the right side than male subjects do.

### A) Mean Length of Tracts

Tract mean length (mm) the average absolute deviation of a data from a central point. We are analysing tract mean length (mm) present in both the sexes (i) Male subjects and ii) Female subjects. The comparison of Mean Length of Fibers is was done between the seed and the end region under four parameters;

a. Male Right and Left Side
b. Female Right and Left side
c. Right side Male and Female
d. Left side Male and Female

**a) Mean Length of Tracts in Male Subjects on both right and left side**

We selected sixteen healthy adult male participants for our study, having a mean age of 30.4 years.

***Right Side:***

The Mean Length of Tracts were analyzed in 16 male subjects by tracing the neural structural connectivity between the primary visual cortex to Inferior Temporal Lobe, we found that the male subject (1008) with the mean age of 57 years having Greater Mean Length of Tracts.

***Left Side:***

Mean Length of Tracts were analyzed in 16 male subjects by tracing the neural structural connectivity between the primary visual cortex to Inferior Temporal Lobe, we found that the male subject (1007) with the mean age of 57 having least Mean Length of Tracts, (Table 9).

**TABLE 9.** MEAN LENGTH OF TRACTS MALE RIGHT AND LEFT*

| Subjects | SIDE | Mean length of Tracts (mm) | SIDE | Mean length of tracts (mm) |
| --- | --- | --- | --- | --- |
| 1006 | R | 69.0259 | L | 78.0762 |
| 1007 | R | 86.0321 | L | 51.0098 |
| 1008 | R | 139.547 | L | 100.312 |
| 1009 | R | 103.704 | L | 91.8397 |
| 1011 | R | 119.8 | L | 107.296 |
| 1013 | R | 89.2041 | L | 111.679 |
| 1014 | R | 96.8163 | L | 40.6223 |
| 1015 | R | 108.12 | L | 74.1402 |
| 1016 | R | 104.525 | L | 76.5809 |
| 1022 | R | 107.692 | L | 62.3265 |
| 1025 | R | 90.9534 | L | 79.3537 |
| 1027 | R | 105.144 | L | 105.24 |
| 1028 | R | 80.9345 | L | 82.9008 |
| 1029 | R | 94.4569 | L | 94.1059 |
| 1030 | R | 106.859 | L | 100.927 |
| 1031 | R | 112.151 | L | 53.6572 |

The $p$ -value is .008685. \*\*The result was statistically significant at $p < .10$

**Fig 14.**
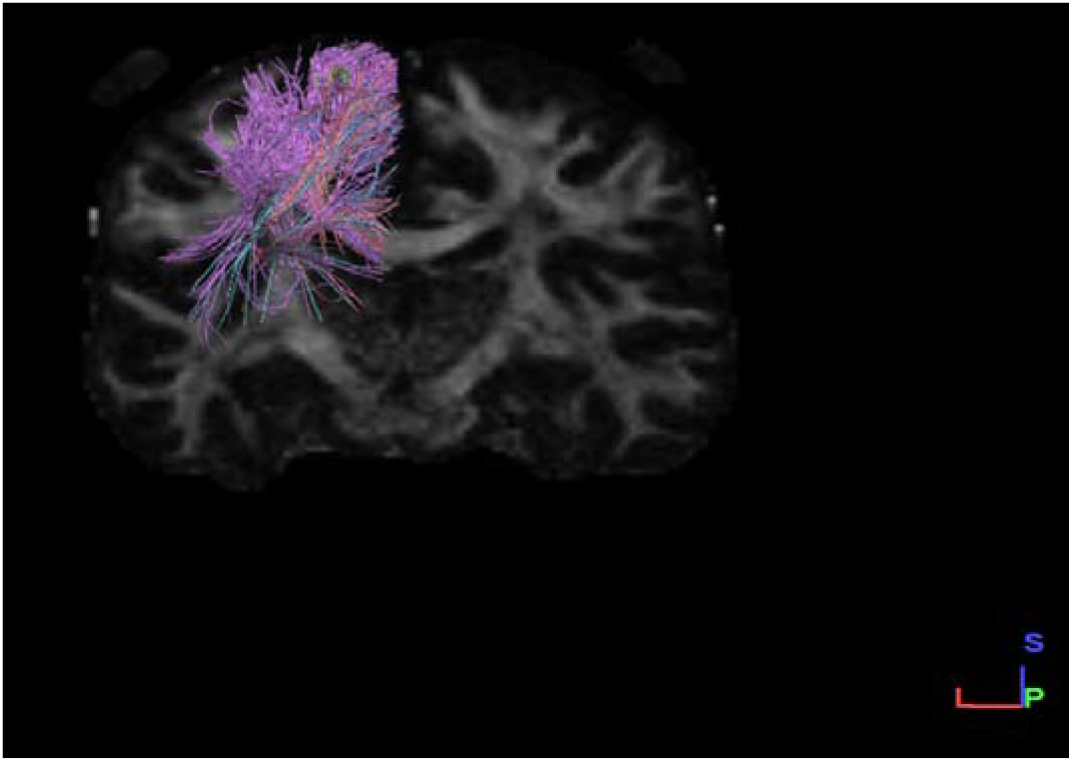
Coronal section male left side dataset

**Fig 15.**
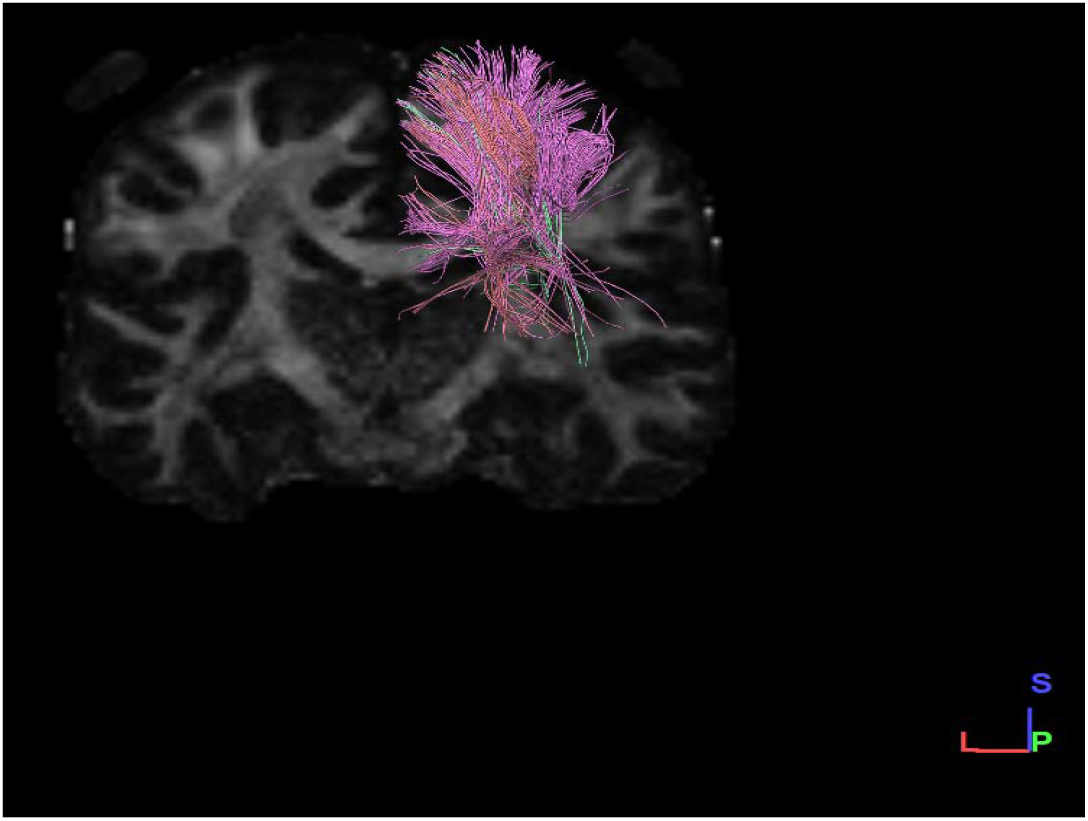
Coronal section male right side dataset

**<u>Graphical Representation</u>**

**Graph-8.**
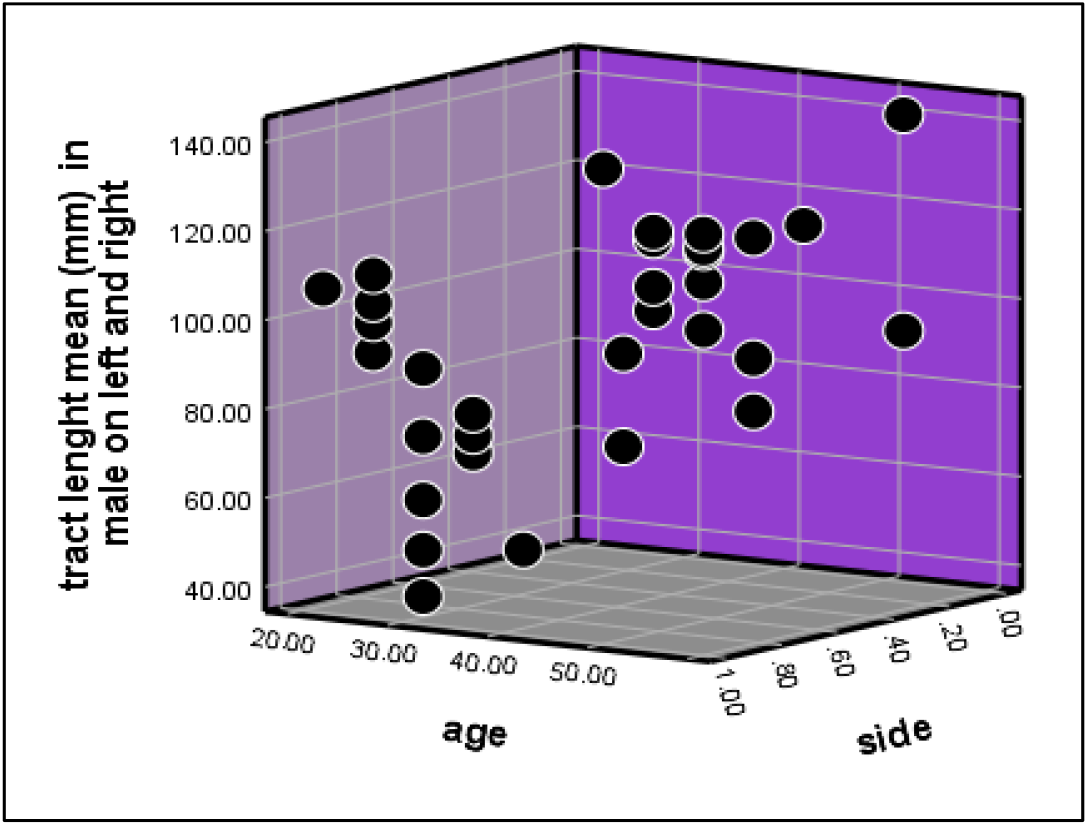
Graphical representation of mean tract length in on both right and left sides in male subjects.

We found that the right side has greater mean tract length than left side in male subjects.

**b) Mean Length of Tracts in Female Subjects on both right and left side**

We selected sixteen healthy adult female participants for our study having a mean age of 30.4 years.

***Right Side:***

The mean length of tracts were analyzed in 16 female subjects by tracing the neural structural connectivity between the primary visual cortex to Inferior Temporal Lobe, we found that the female subject (1004) with the mean age of 45 years having greater mean length of tracts.

***Left Side:***

Mean length of tracts were analyzed in 16 female subjects by tracing the neural structural connectivity between the primary visual cortex to Inferior Temporal Lobe, we found that the female subject (1001) with the mean age of 41.4 having least mean length of tracts, (Table 10).

**TABLE 10.**
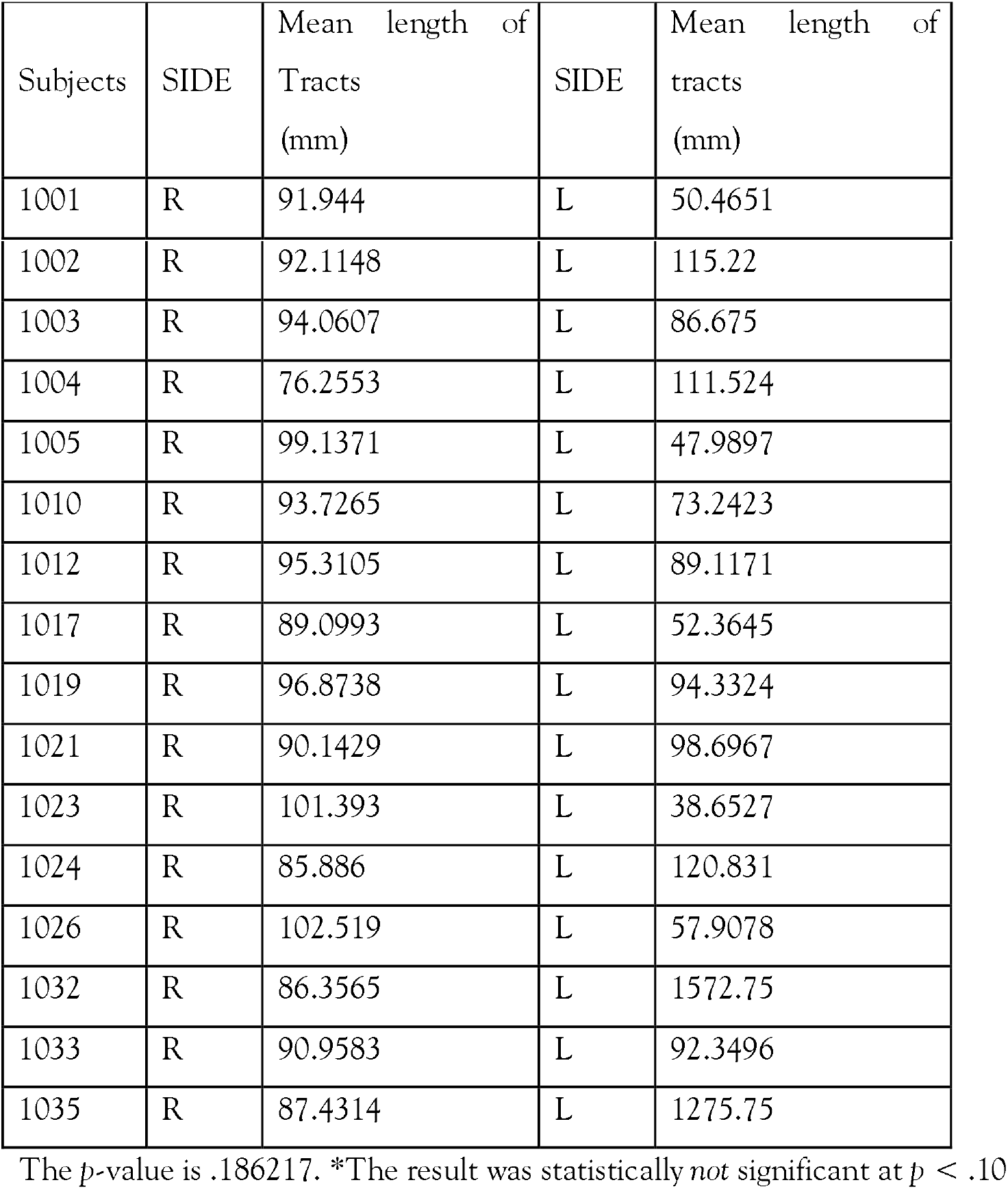
MEAN LENGTH OF TRACTS IN FEMALE RIGHT AND LEFT*

| Subjects | SIDE | Mean length of Tracts (mm) | SIDE | Mean length of tracts (mm) |
| --- | --- | --- | --- | --- |
| 1001 | R | 91.944 | L | 50.4651 |
| 1002 | R | 92.1148 | L | 115.22 |
| 1003 | R | 94.0607 | L | 86.675 |
| 1004 | R | 76.2553 | L | 111.524 |
| 1005 | R | 99.1371 | L | 47.9897 |
| 1010 | R | 93.7265 | L | 73.2423 |
| 1012 | R | 95.3105 | L | 89.1171 |
| 1017 | R | 89.0993 | L | 52.3645 |
| 1019 | R | 96.8738 | L | 94.3324 |
| 1021 | R | 90.1429 | L | 98.6967 |
| 1023 | R | 101.393 | L | 38.6527 |
| 1024 | R | 85.886 | L | 120.831 |
| 1026 | R | 102.519 | L | 57.9078 |
| 1032 | R | 86.3565 | L | 1572.75 |
| 1033 | R | 90.9583 | L | 92.3496 |
| 1035 | R | 87.4314 | L | 1275.75 |

The $p$ -value is .186217. \*The result was statistically *not* significant at $p < .10$

**Fig 16.**
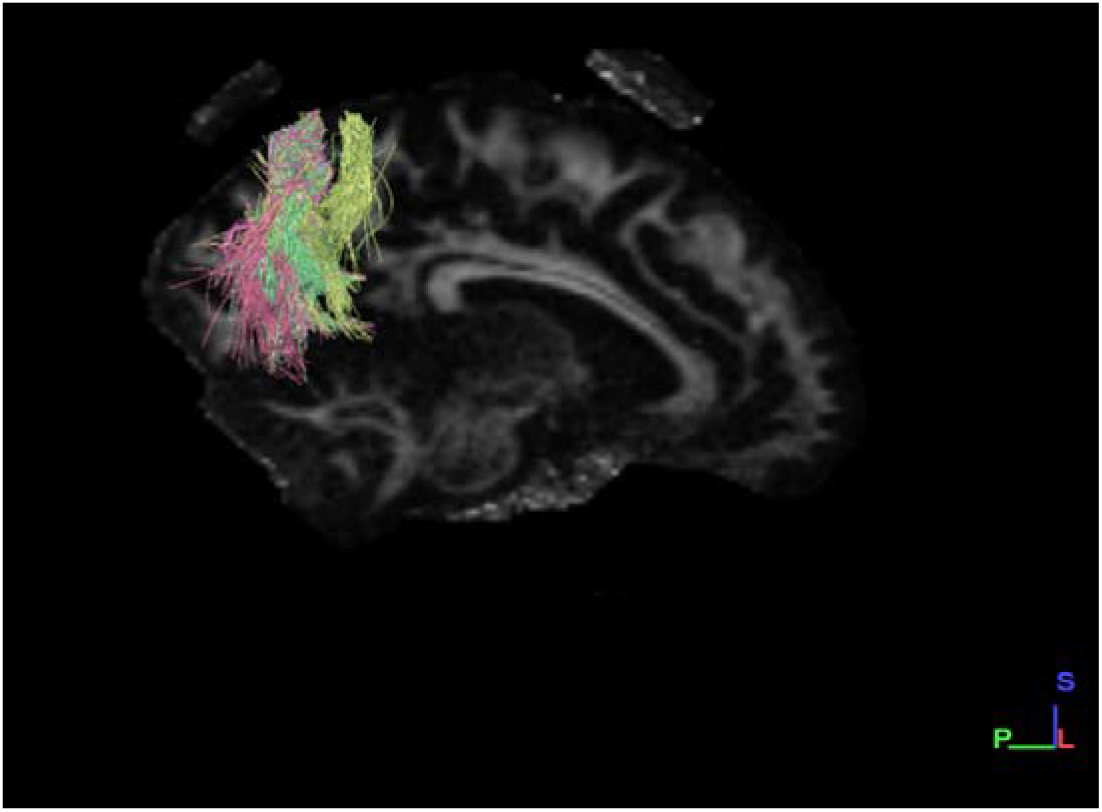
Sagittal section female right side dataset

**Fig 17.**
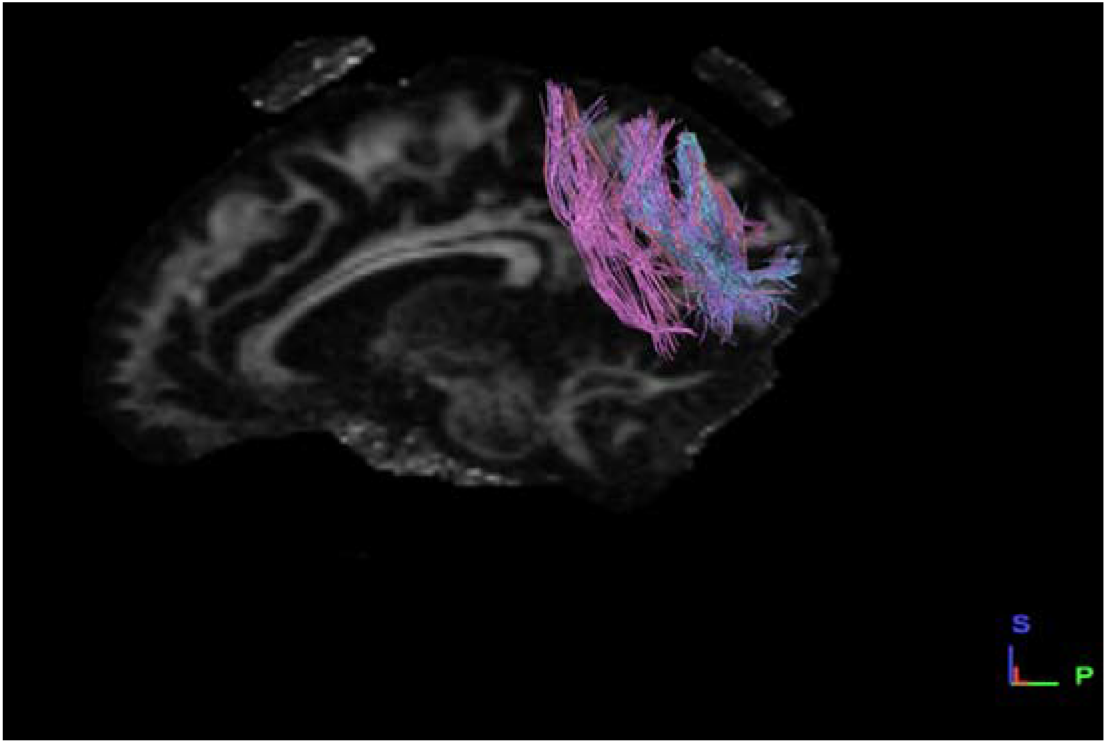
Sagittal section female left side dataset

**<u>Graphical Representation</u>**

**Graph-9.**
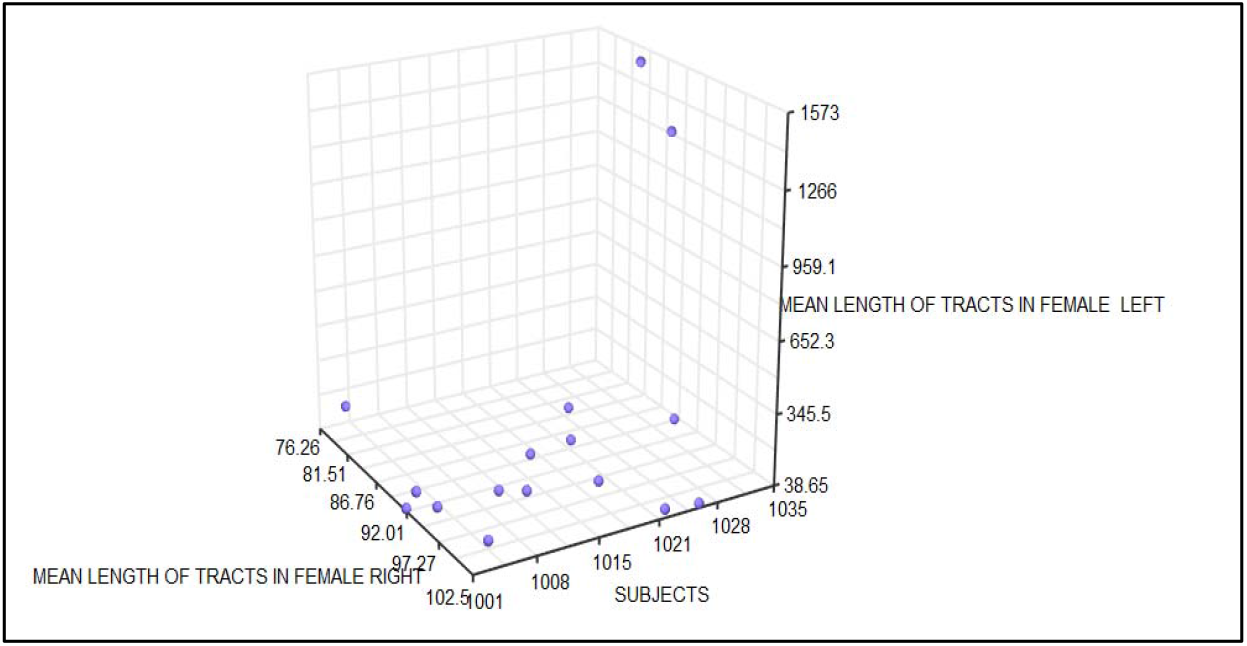
Graphical representation of mean tract length in on both right and left sides in female subjects.

We found that the right side is having greater number of tracts than left side in female subjects.

**c) Mean Length of Tracts both Male and Female Left Side**

We selected sixteen healthy adult male and female participants for our study having a mean age of 30.4 years.

***Male:***

The mean length of tracts were analyzed in 16 male subjects by tracing the neural structural connectivity between the primary visual cortex to Inferior Temporal Lobe, we found that the male subject (1008) with the mean age of 57 years having greater mean length of tracts.

***Female:***

Mean Length of Tracts were analyzed in 16 female subjects by tracing the neural structural connectivity between the primary visual cortex to Inferior Temporal Lobe, we found that the female subject (1004) with the mean age of 27 having least mean length of tracts, (Table 11).

**TABLE 11.** MEAN LENGTH OF TRACTS MALE AND FEMALE RIGHT**

| Female<br>Datasets | Mean length of Tracts<br>(mm) | Male<br>Datasets | Mean length of tracts<br>(mm) |
| --- | --- | --- | --- |
| 1001 | 91.944 | 1006 | 69.0259 |
| 1002 | 92.1148 | 1007 | 86.0321 |
| 1003 | 94.0607 | 1008 | 139.547 |
| 1004 | 76.2553 | 1009 | 103.704 |
| 1005 | 99.1371 | 1011 | 119.8 |
| 1010 | 93.7265 | 1013 | 89.2041 |
| 1012 | 95.3105 | 1014 | 96.8163 |
| 1017 | 89.0993 | 1015 | 108.12 |
| 1019 | 96.8738 | 1016 | 104.525 |
| 1021 | 90.1429 | 1022 | 107.692 |
| 1023 | 101.393 | 1025 | 90.9534 |
| 1024 | 85.886 | 1027 | 105.144 |
| 1026 | 102.519 | 1028 | 80.9345 |
| 1032 | 86.3565 | 1029 | 94.4569 |
| 1033 | 90.9583 | 1030 | 106.859 |
| 1035 | 87.4314 | 1031 | 112.151 |

The $p$ -value is .05493. \*\*The result was statistically significant at $p < .10$

**Fig 18.**
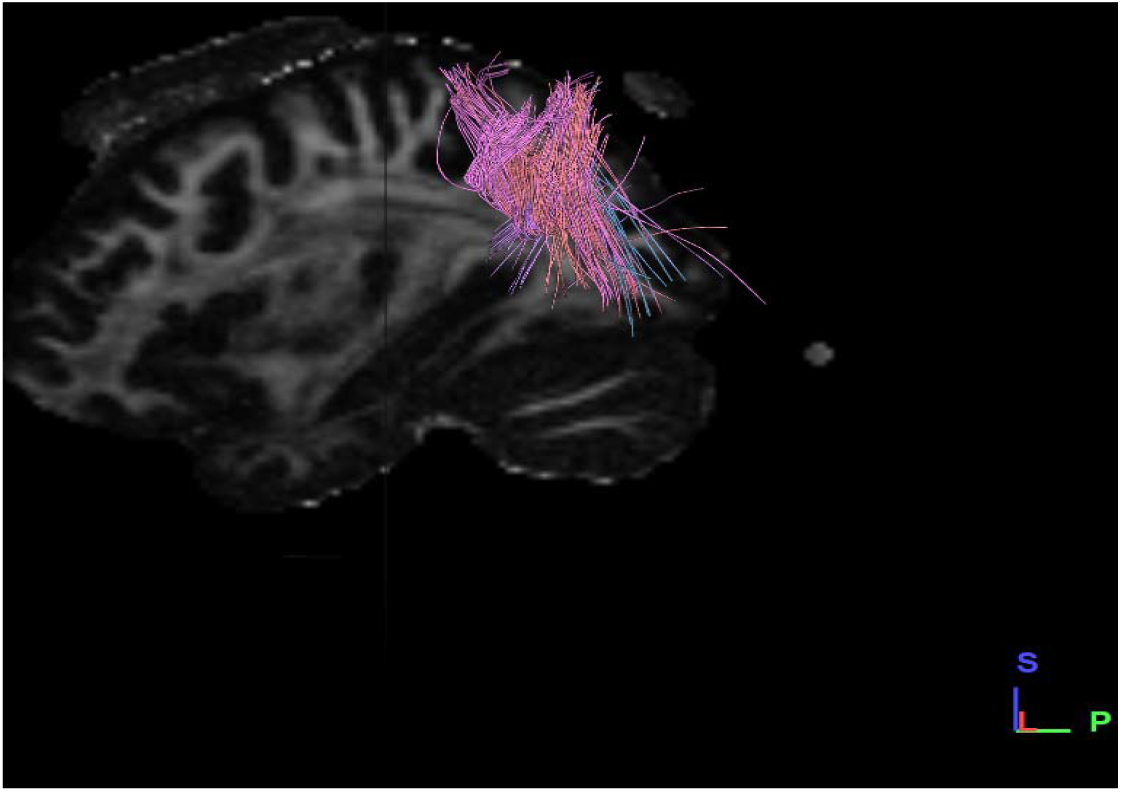
Sagittal section female right side dataset

**Fig 19.**
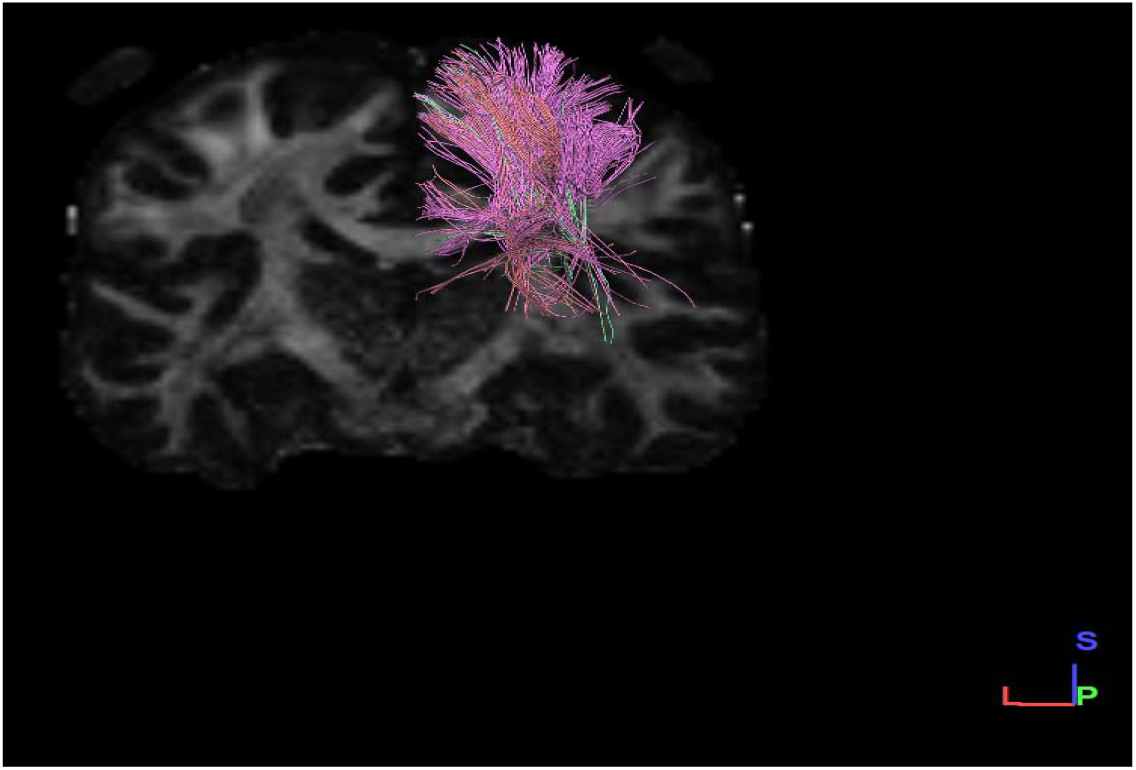
Coronal section male right side dataset

**<u>Graphical Representation</u>**

**Graph-10.**
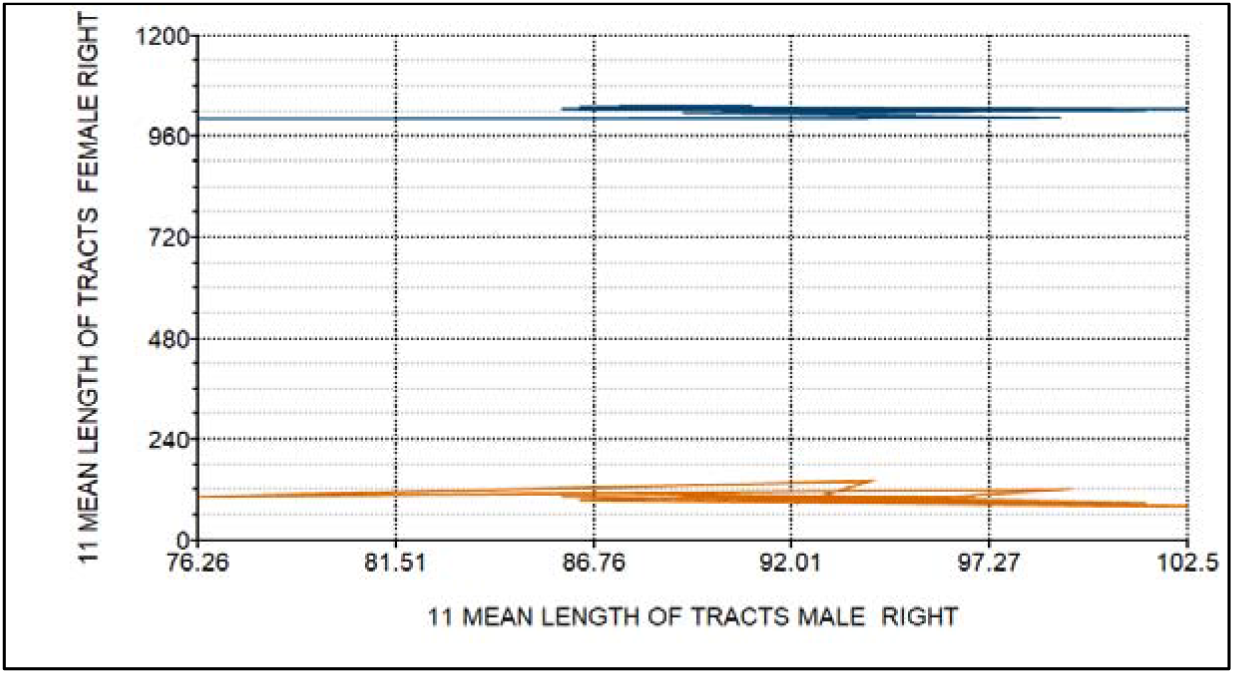
Graphical representation of mean tract length in both male and female subjects on right side.

We found that the male subjects have greater mean length of tract fibres on right side than female subjects.

**d) Mean Length of Tracts both Male and Female Right Side**

We selected sixteen healthy adults male and female participants for our study having a mean age of 30.4 years.

***Male:***

The Mean Length of Tracts were analyzed in 16 male subjects by tracing the neural structural connectivity between the primary visual cortex to Inferior Temporal Lobe, we found that the male subject (1032) with the mean age of 27 years having Greater Mean Length of Tracts.

***Female:***

Mean Length of Tracts were analyzed in 16 female subjects by tracing the neural structural connectivity between the primary visual cortex to Inferior Temporal Lobe, we found that the female subject (1007) with the mean age of 27 having least Mean Length of Tracts, (Table 12).

**TABLE 12.** MEAN LENGTH OF TRACTS MALE AND FEMALE LEFT*

| Female Datasets | Mean length of Tracts (mm) | Male Datasets | Mean length of tracts (mm) |
| --- | --- | --- | --- |
| 1001 | 50.4651 | 1006 | 78.0762 |
| 1002 | 115.22 | 1007 | 51.0098 |
| 1003 | 86.675 | 1008 | 100.312 |
| 1004 | 111.524 | 1009 | 91.8397 |
| 1005 | 47.9897 | 1011 | 107.296 |
| 1010 | 73.2423 | 1013 | 111.679 |
| 1012 | 89.1171 | 1014 | 40.6223 |
| 1017 | 52.3645 | 1015 | 74.1402 |
| 1019 | 94.3324 | 1016 | 76.5809 |
| 1021 | 98.6967 | 1022 | 62.3265 |
| 1023 | 38.6527 | 1025 | 79.3537 |
| 1024 | 120.831 | 1027 | 105.24 |
| 1026 | 57.9078 | 1028 | 82.9008 |
| 1032 | 1572.75 | 1029 | 94.1059 |
| 1033 | 92.3496 | 1030 | 100.927 |
| 1035 | 1275.75 | 1031 | 89.0411 |

The $p$ -value is .165737. \*The result was statistically *not* significant at $p < .10$ .

**Fig 20.**
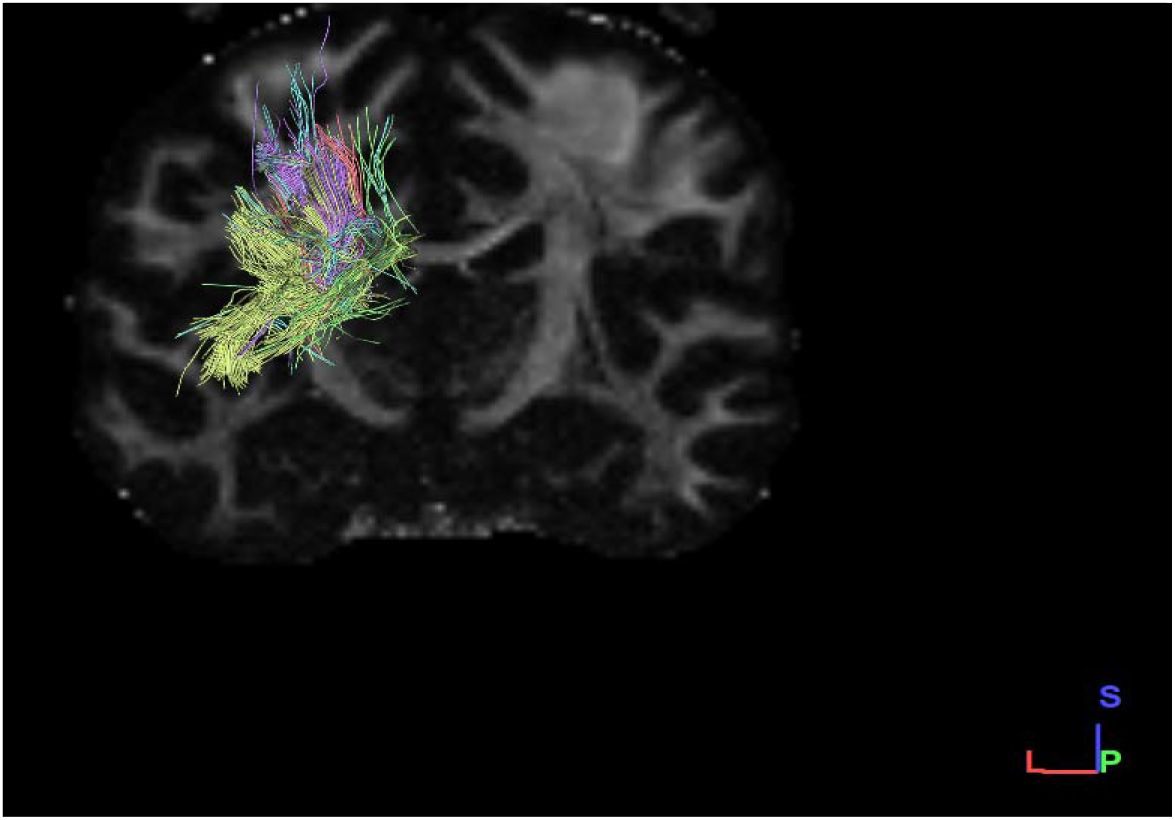
Coronal section male left side dataset

**Fig 21.**
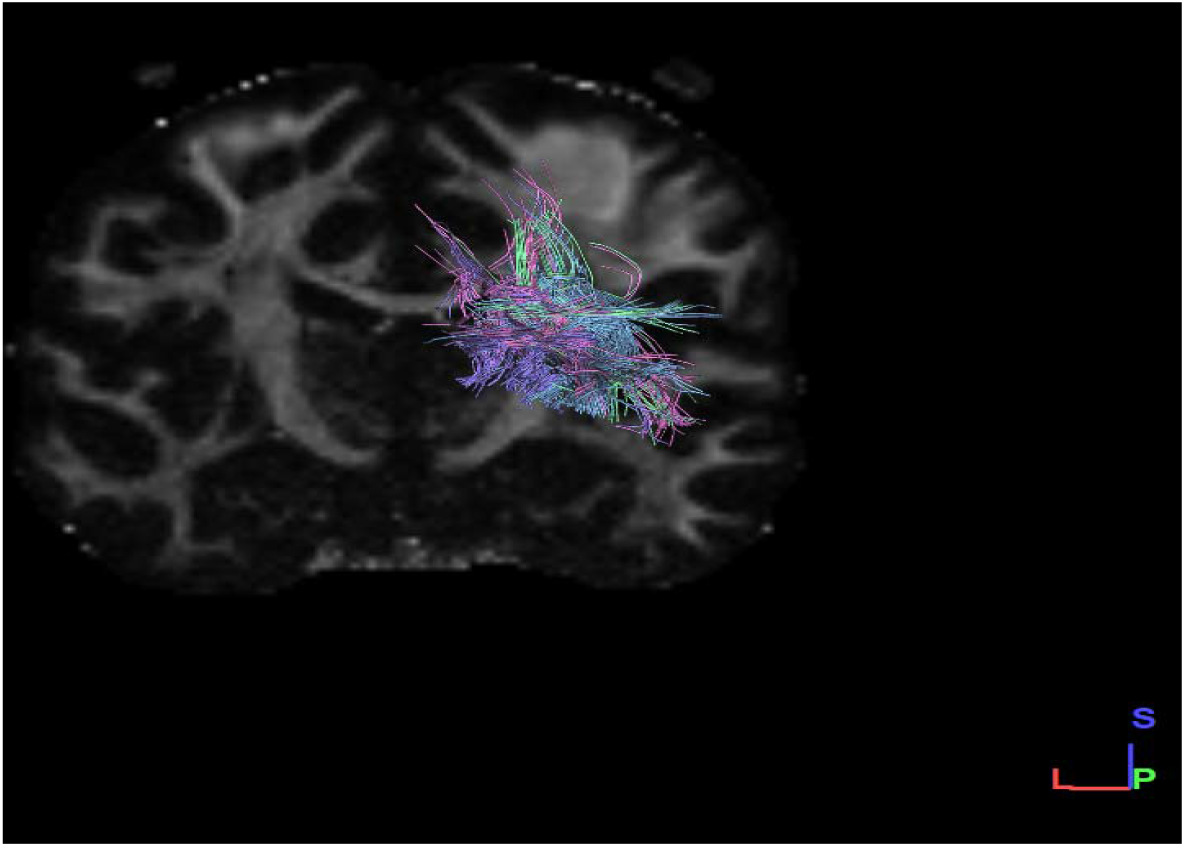
Coronal section Female left side dataset

**<u>Graphical Representation</u>**

**Grap-11.**
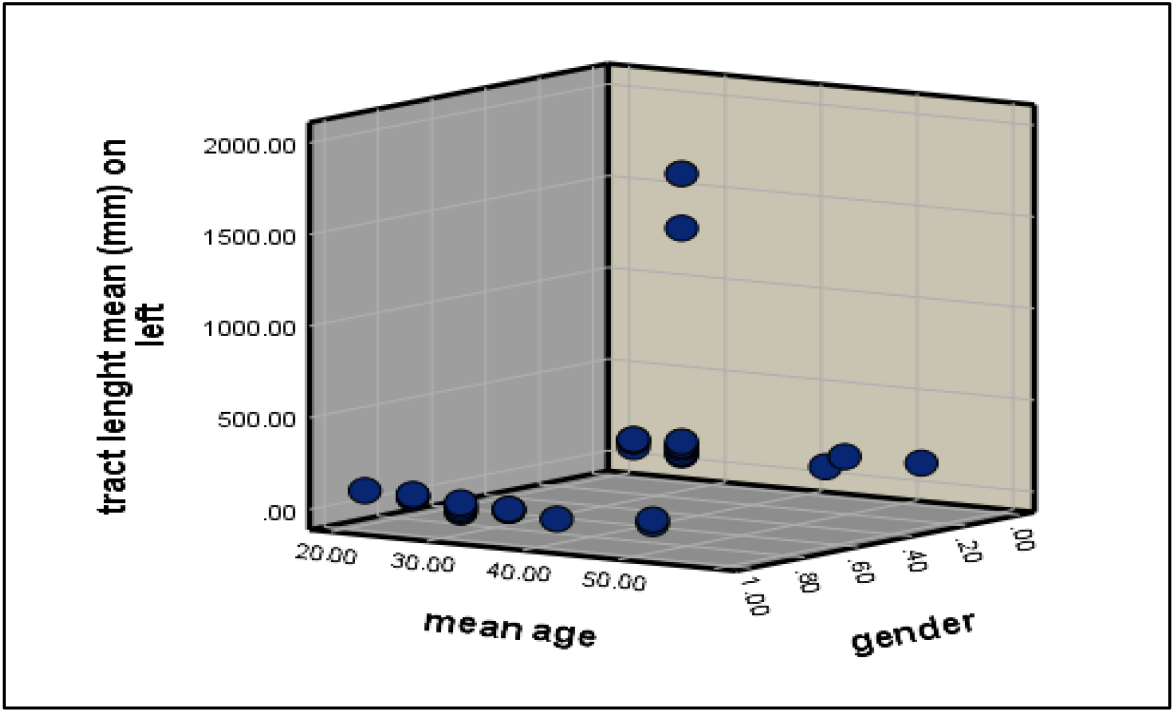
Graphical representation of mean tract length in both male and female subjects on left side.

We found that the male subjects have greater mean tract length of fibers on left side than female subjects.

### D) Tracts Length Standard Deviation

**a) Tract length standard deviation in male subjects on right and left side**

We selected sixteen healthy adult male participants for our study, having a mean age of 30.4 years.

***Right Side:***

The Tract length standard deviation were analyzed in 16 male subjects by tracing the neural structural connectivity between the primary visual cortex to Inferior Temporal Lobe, we found that the male subject (1009) with the mean age of years has least tract length standard deviation.

***Left Side:***

Tract length standard deviation were analyzed in 16 male subjects by tracing the neural structural connectivity between the primary visual cortex to Inferior Temporal Lobe, we found that the male subject (1029) with the mean age of having highest tract length standard deviation, (Table 13).

**TABLE 13.**
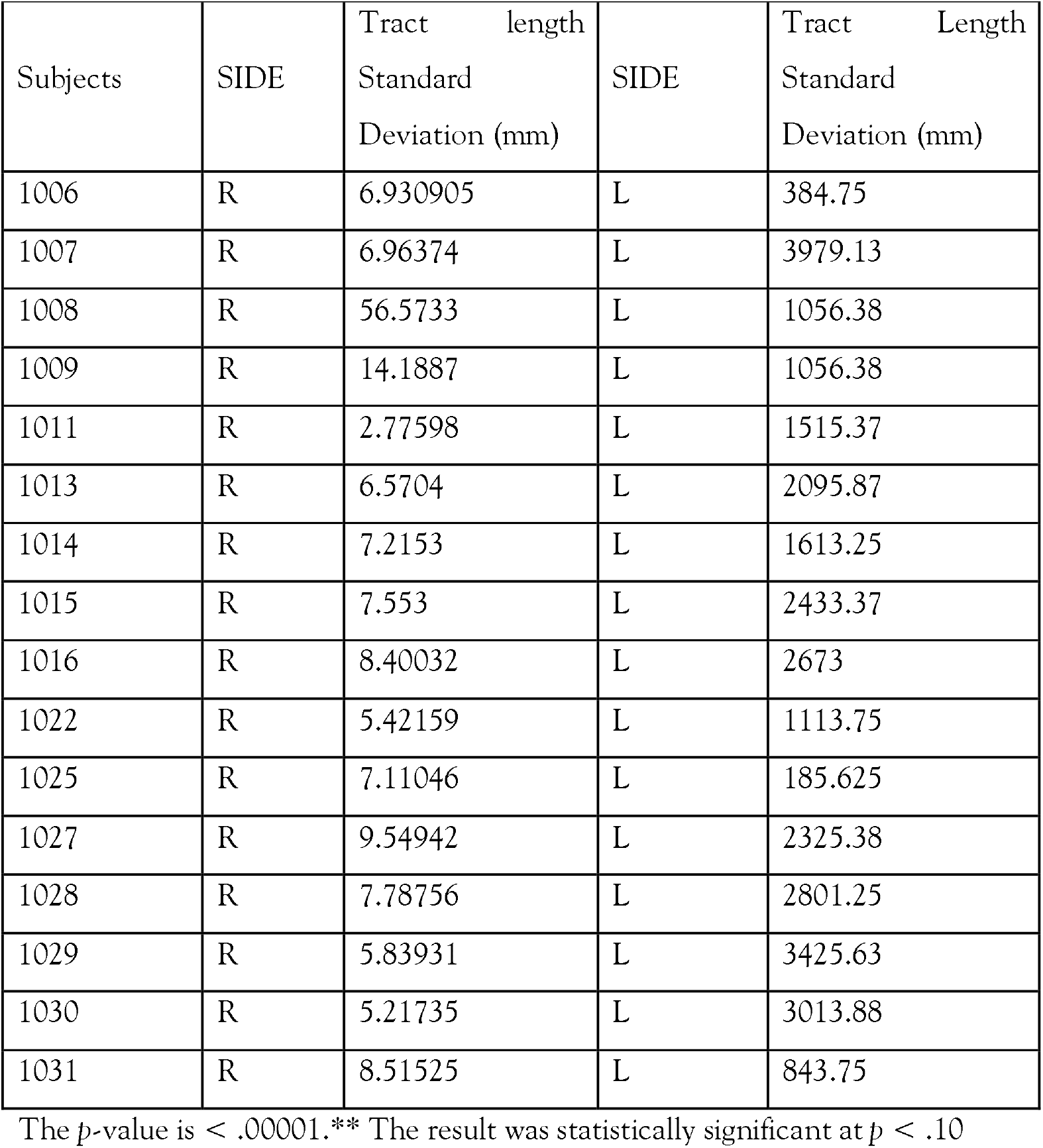
TRACT LENGTH STANDARD DEVIATION IN MALE RIGHT AND LEFT**

| Subjects | SIDE | Tract length<br>Standard<br>Deviation (mm) | SIDE | Tract Length<br>Standard<br>Deviation (mm) |
| --- | --- | --- | --- | --- |
| 1006 | R | 6.930905 | L | 384.75 |
| 1007 | R | 6.96374 | L | 3979.13 |
| 1008 | R | 56.5733 | L | 1056.38 |
| 1009 | R | 14.1887 | L | 1056.38 |
| 1011 | R | 2.77598 | L | 1515.37 |
| 1013 | R | 6.5704 | L | 2095.87 |
| 1014 | R | 7.2153 | L | 1613.25 |
| 1015 | R | 7.553 | L | 2433.37 |
| 1016 | R | 8.40032 | L | 2673 |
| 1022 | R | 5.42159 | L | 1113.75 |
| 1025 | R | 7.11046 | L | 185.625 |
| 1027 | R | 9.54942 | L | 2325.38 |
| 1028 | R | 7.78756 | L | 2801.25 |
| 1029 | R | 5.83931 | L | 3425.63 |
| 1030 | R | 5.21735 | L | 3013.88 |
| 1031 | R | 8.51525 | L | 843.75 |

The $p$ -value is $< .00001$ .\*\* The result was statistically significant at $p < .10$

**Fig 22.**
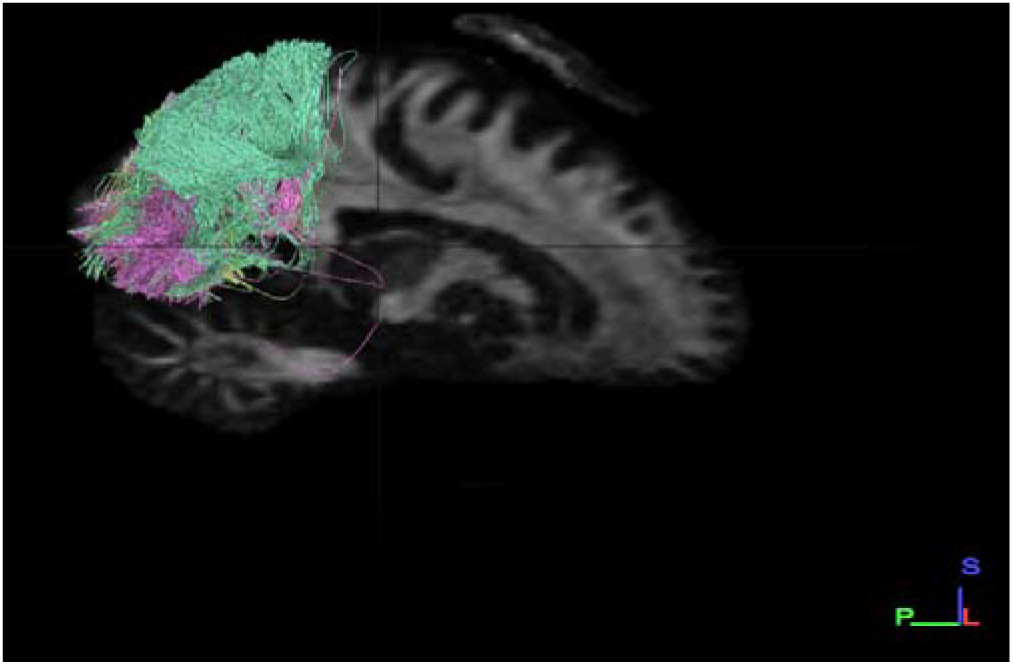
Sagittal section male right side dataset

**Fig 23.**
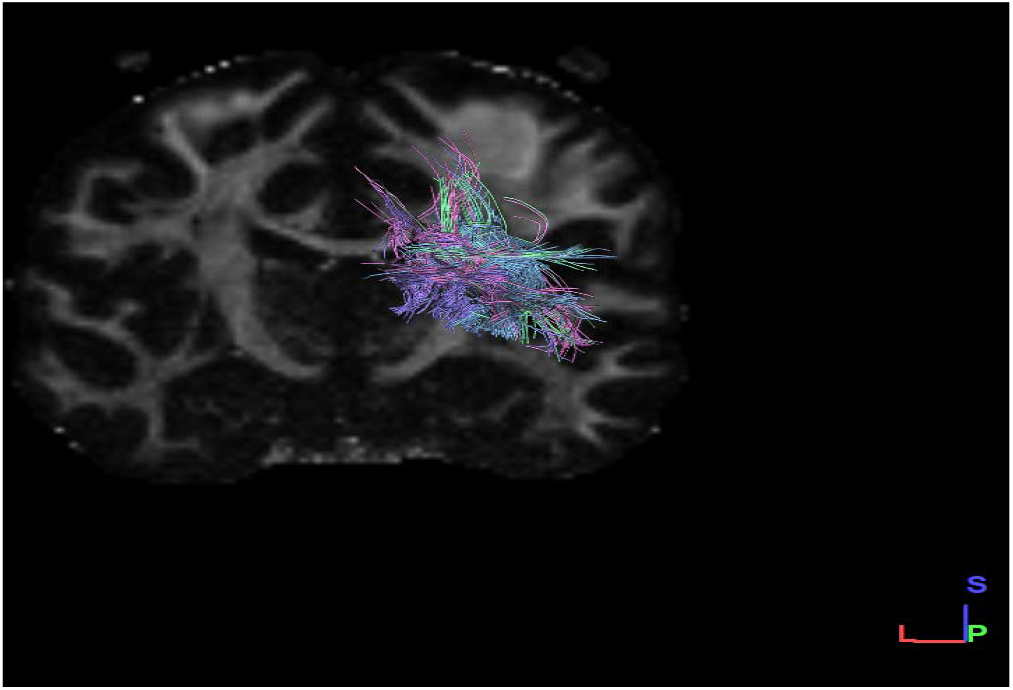
Coronal section male left side dataset

**<u>Graphical Representation</u>**

**Graph-12.**
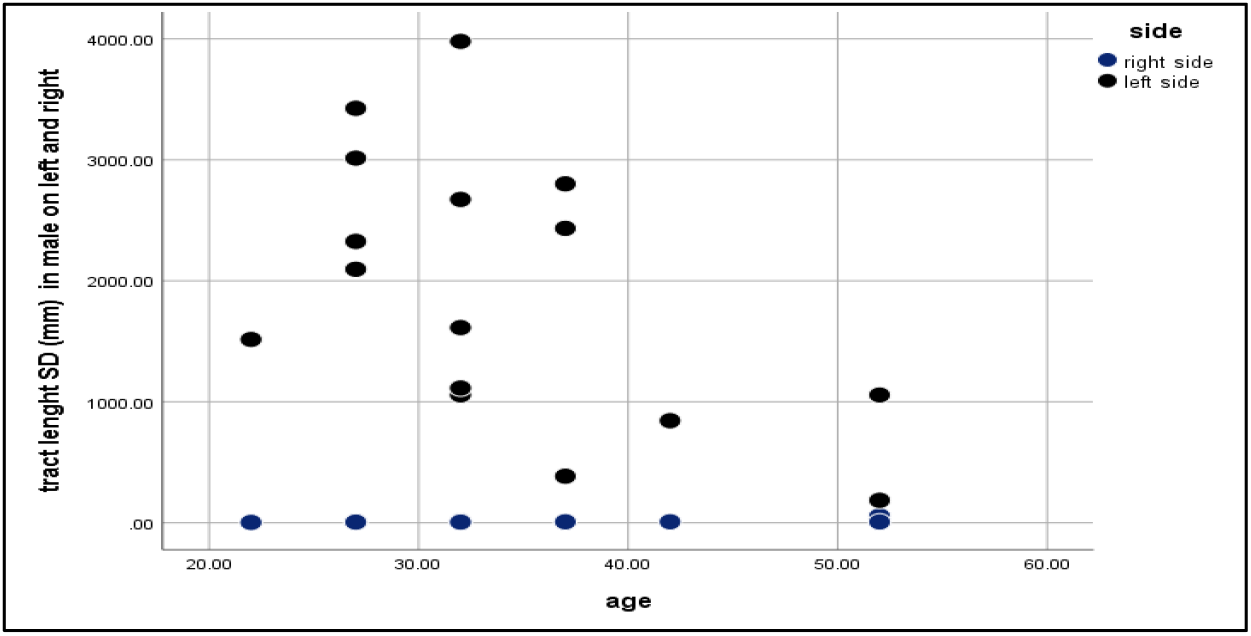
Graphical representation of tract length standard deviation on both right and left sides in male subjects.

We found that male subjects have greater tracts length standard deviation on left side than on right side.

**b) Tract Length Standard Deviation in Female subjects on both Right and Left Sides**

We selected sixteen healthy adult male participants for our study, having a mean age of 30.4 years.

***Right Side:***

The Tract length standard deviation were analyzed in 16 female subjects by tracing the neural structural connectivity between the primary visual cortex to Inferior Temporal Lobe, we found that the female subject have least tract standard deviation on right side (1007).

***Left Side:***

Tract length standard deviation were analyzed in 16 female subjects by tracing the neural structural connectivity between the primary visual cortex to Inferior Temporal Lobe, we found least tract length standard deviation in female subjects on the right side (1007), (Table 14).

**TABLE 14.** TRACT LENGTH STANDARD DEVIATION IN FEMALE RIGHT AND LEFT**

| Subjects | SIDE | Tract length<br>Standard<br>Deviation (mm) | SIDE | Tract Length<br>Standard Deviation<br>(mm) |
| --- | --- | --- | --- | --- |
| 1001 | R | 7.76203 | L | 118.125 |
| 1002 | R | 7.02543 | L | 8032.5 |
| 1003 | R | 8.47677 | L | 9321.75 |
| 1004 | R | 8.99159 | L | 2956.5 |
| 1005 | R | 7.96865 | L | 1117.12 |
| 1010 | R | 8.73392 | L | 853.875 |
| 1012 | R | 7.25344 | L | 421.875 |
| 1017 | R | 12.0259 | L | 3931.87 |
| 1019 | R | 4.68937 | L | 3894.75 |
| 1021 | R | 4.57601 | L | 4131 |
| 1023 | R | 7.43073 | L | 2953.13 |
| 1024 | R | 6.25829 | L | 1420.88 |
| 1026 | R | 6.32936 | L | 3628.13 |
| 1032 | R | 4.8742 | L | 4.8742 |
| 1033 | R | 9.60303 | L | 1917 |
| 1035 | R | 6.07921 | L | 6.07921 |

The $p$ -value is .000317. \*\*The result was statistically significant at $p < .10$

**Fig 24.**
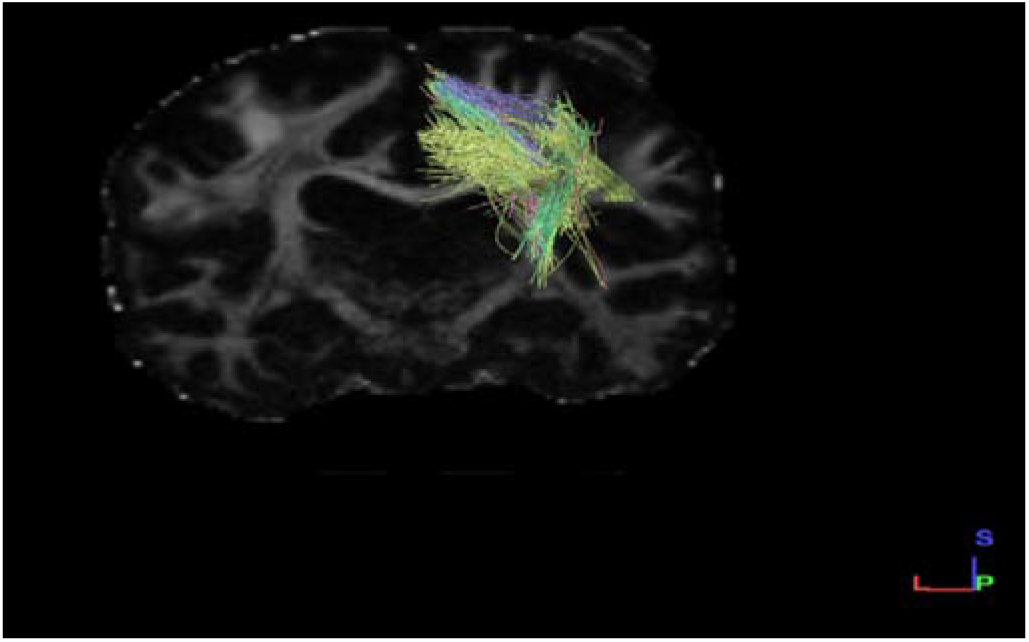
Coronal section Female right side dataset

**Fig25.**
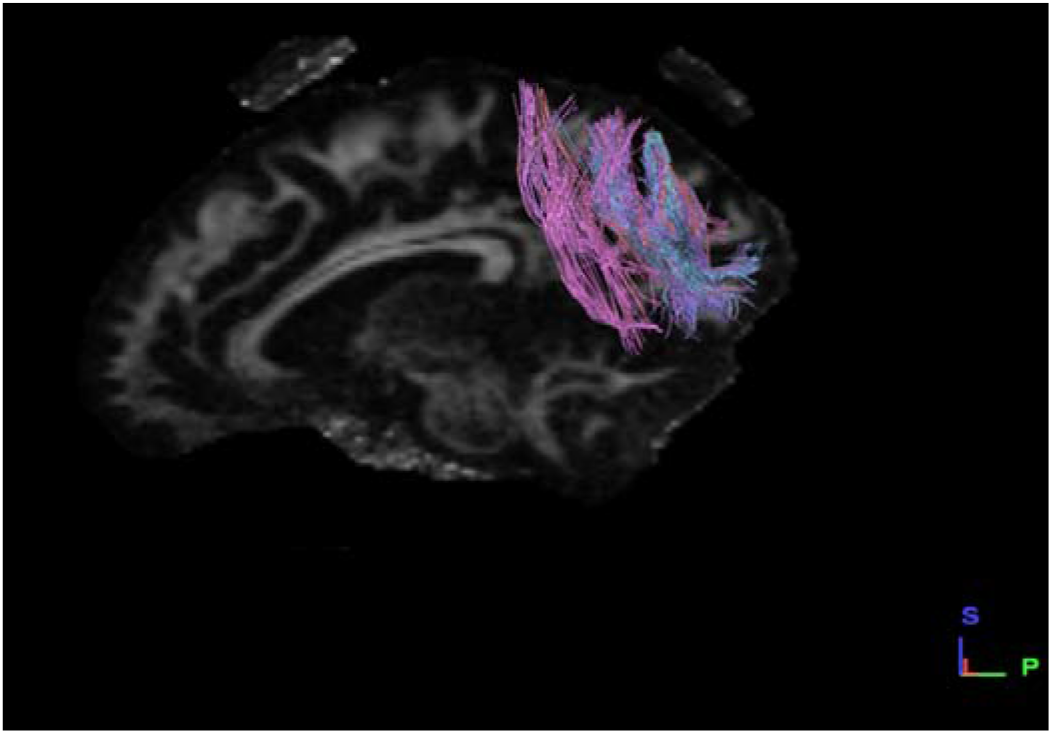
Sagittal section Female left side dataset

**<u>Graphical Representation</u>**

**Graph-13.**
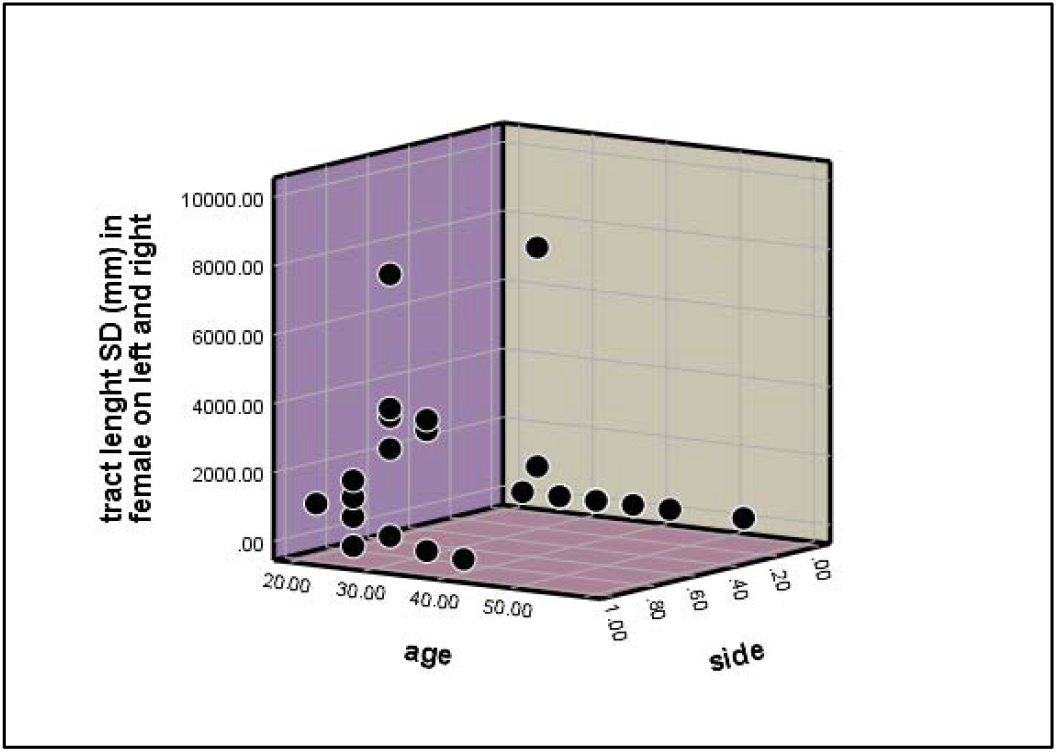
Graphical representation of tract length standard deviation on both right and left sides in female subjects.

We found that left side of female subjects has greater tract length standard deviation than right side.

**c) Tract Length Standard Deviation in both Male and Female Right Side**

***Male:***

The tract length standard deviation was analyzed in 16 male subjects by tracing the neural structural connectivity between the primary visual cortex to Inferior Temporal Lobe, we found that the male subject (1008) with the mean age of 57 years has greater mean length of tracts.

***Female:***

The tracts length standard deviation were analyzed in 16 female subjects by tracing the neural structural connectivity between the primary visual cortex to Inferior Temporal Lobe, we found that the female subject (1021) with the mean age of 27 having least mean length of tracts, (Table 9).

**TABLE 15.** TRACT LENGTH STANDARD DEVIATION MALE AND FEMALE AT RIGHT*

| Female datasets | Tract length Standard Deviation (mm) | Male Datasets | Tract Length Standard Deviation (mm) |
| --- | --- | --- | --- |
| 1001 | 7.76203 | 1006 | 6.930905 |
| 1002 | 7.02543 | 1007 | 6.96374 |
| 1003 | 8.47677 | 1008 | 56.5733 |
| 1004 | 8.99159 | 1009 | 14.1887 |
| 1005 | 7.96865 | 1011 | 2.77598 |
| 1010 | 8.73392 | 1013 | 6.5704 |
| 1012 | 7.25344 | 1014 | 7.2153 |
| 1017 | 12.0259 | 1015 | 7.553 |
| 1019 | 4.68937 | 1016 | 8.40032 |
| 1021 | 4.57601 | 1022 | 5.42159 |
| 1023 | 7.43073 | 1025 | 7.11046 |
| 1024 | 6.25829 | 1027 | 9.54942 |
| 1026 | 6.32936 | 1028 | 7.78756 |
| 1032 | 4.8742 | 1029 | 5.83931 |
| 1033 | 9.60303 | 1030 | 5.21735 |
| 1035 | 6.07921 | 1031 | 8.51525 |

The $p$ -value is .346856. \*The result was statistically *not* significant at $p < .10$

**Fig 26.**
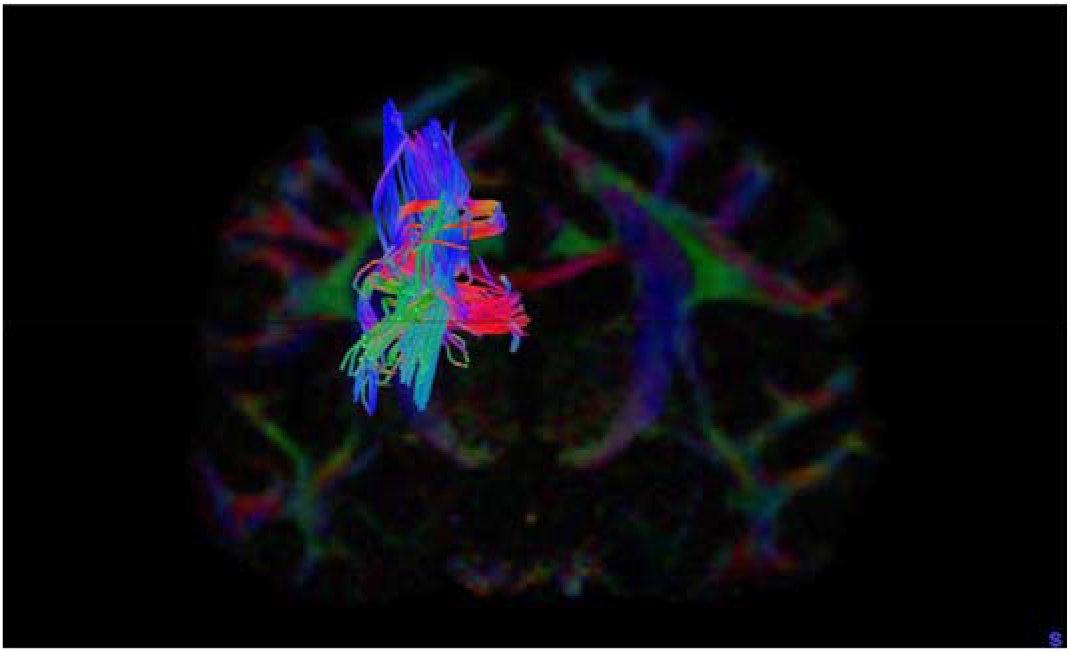
Coronal section male left side dataset

**Fig 27.**
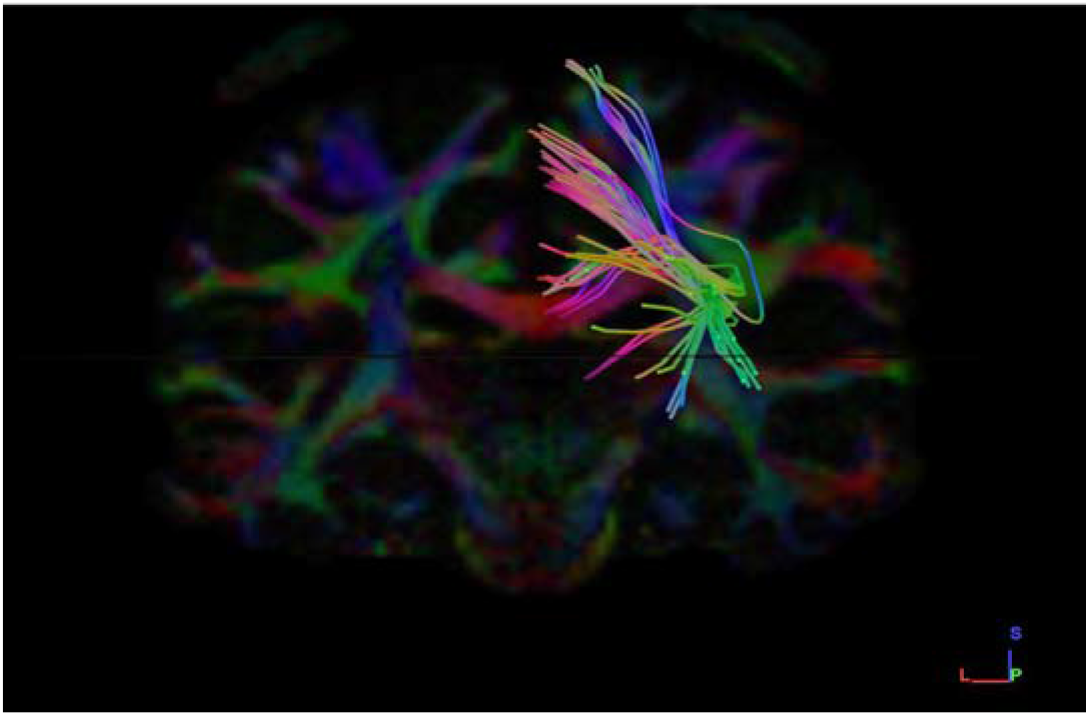
Coronal section female right side dataset

**<u>Graphical Representation</u>**

**Graph-14.**
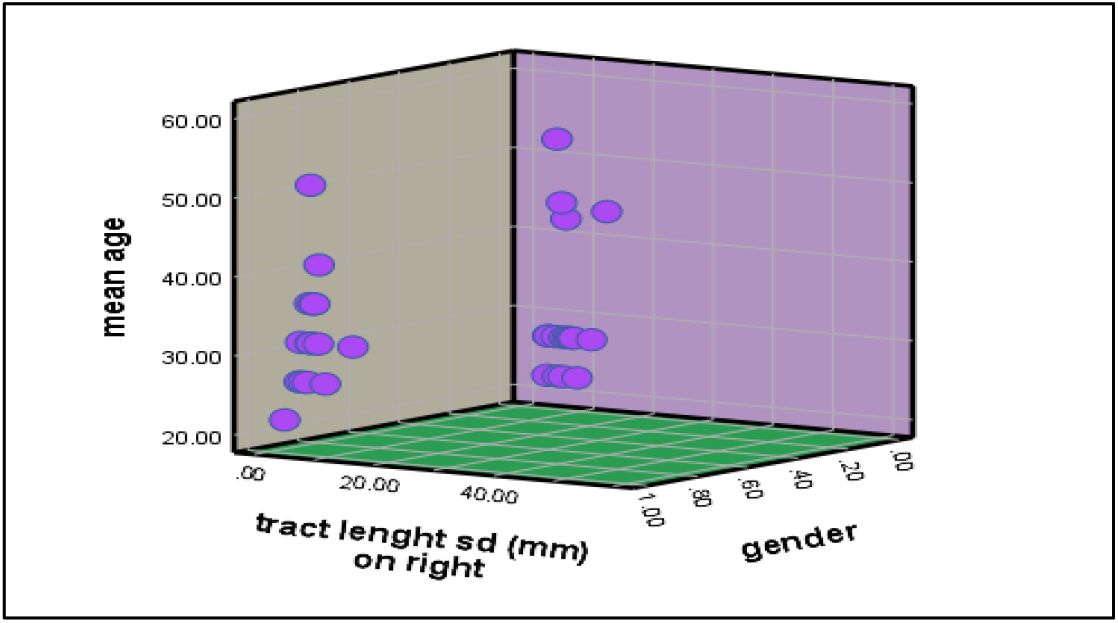
Graphical representation of tract length standard deviation on right side in male and female subjects.

We found that male subjects have greater tract length standard deviation on right side than female subjects.

**d) The Tract Length Standard Deviation both Male and Female Left Side**

***Male:***

The tract length standard deviation was analyzed in 16 male subjects by tracing the neural structural connectivity between the primary visual cortex to Inferior Temporal Lobe, we found that the male subject (1001) with the mean age of 42 years having least Mean Length of Tracts.

***Female:***

Tract length standard deviation were analyzed in 16 female subjects by tracing the neural structural connectivity between the primary visual cortex to Inferior Temporal Lobe, we found that the female subject (1003) with the mean age of 27 has greater Mean Length of Tracts, (Table 9).

**TABLE 16.** TRACT LENGTH STANDARD DEVIATION IN MALE AND FEMALE AT LEFT *

| Female datasets | Tract length Standard Deviation (mm) | Male Datasets | Tract Length Standard Deviation (mm) |
| --- | --- | --- | --- |
| 1001 | 118.125 | 1006 | 384.75 |
| 1002 | 8032.5 | 1007 | 3979.13 |
| 1003 | 9321.75 | 1008 | 1056.38 |
| 1004 | 2956.5 | 1009 | 1056.38 |
| 1005 | 1117.12 | 1011 | 1515.37 |
| 1010 | 853.875 | 1013 | 2095.87 |
| 1012 | 421.875 | 1014 | 1613.25 |
| 1017 | 3931.87 | 1015 | 2433.37 |
| 1019 | 3894.75 | 1016 | 2673 |
| 1021 | 4131 | 1022 | 1113.75 |
| 1023 | 2953.13 | 1025 | 185.625 |
| 1024 | 1420.88 | 1027 | 2325.38 |
| 1026 | 3628.13 | 1028 | 2801.25 |
| 1032 | 4.8742 | 1029 | 3425.63 |
| 1033 | 1917 | 1030 | 3013.88 |
| 1035 | 6.07921 | 1031 | 843.75 |

The $p$ -value is .239442. \*The result was statistically *not* significant at $p < .10$ .

**Fig 28.**
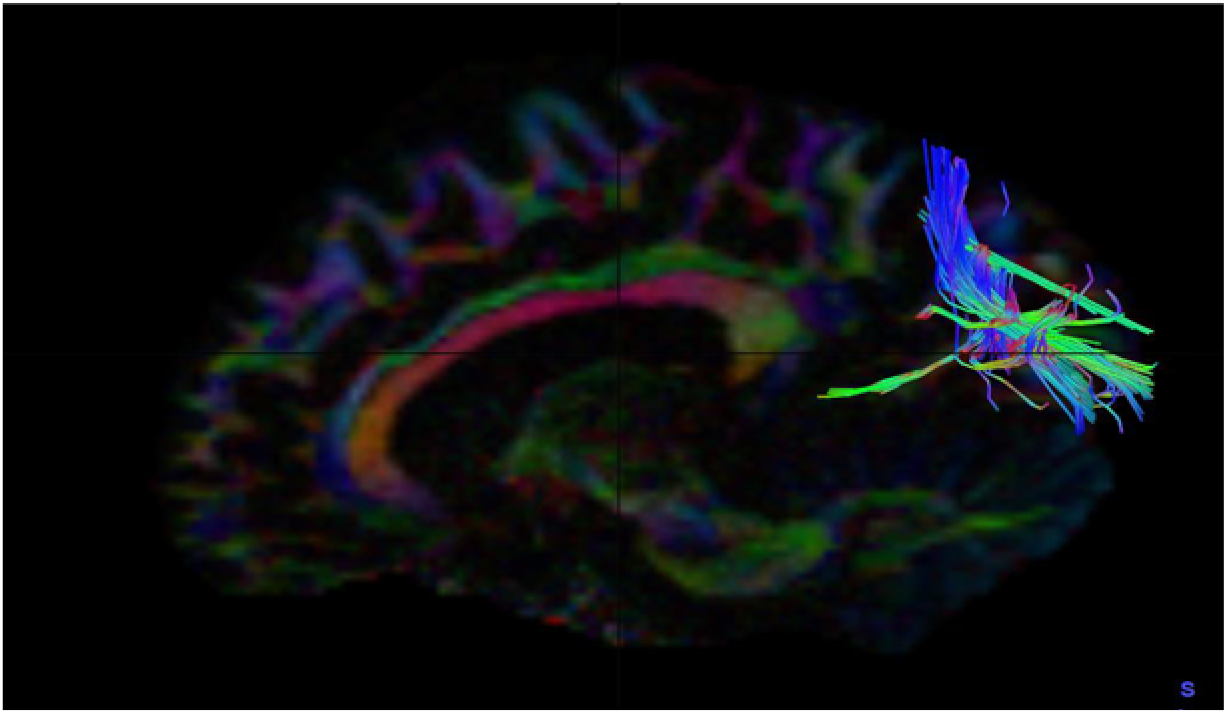
Sagittal section male left side dataset

**Fig 29.**
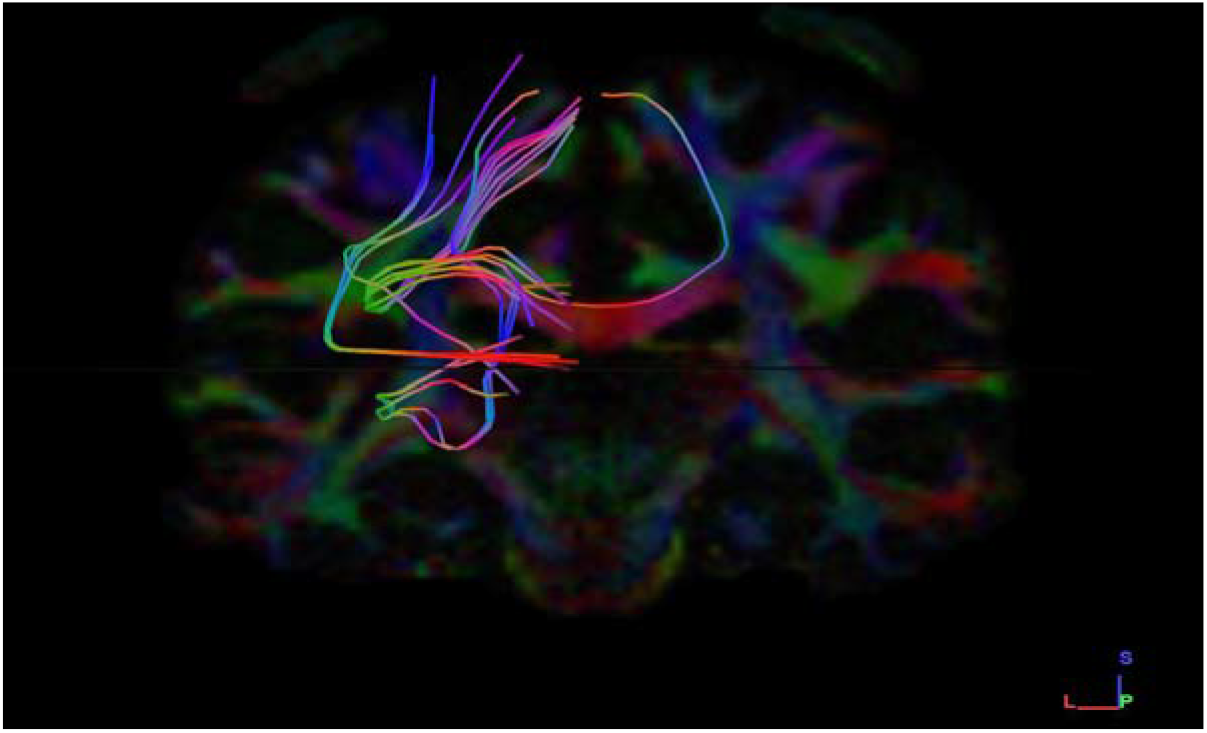
Coronal section Female left side dataset

**<u>Graphical Representation</u>**

**Graph-15.**
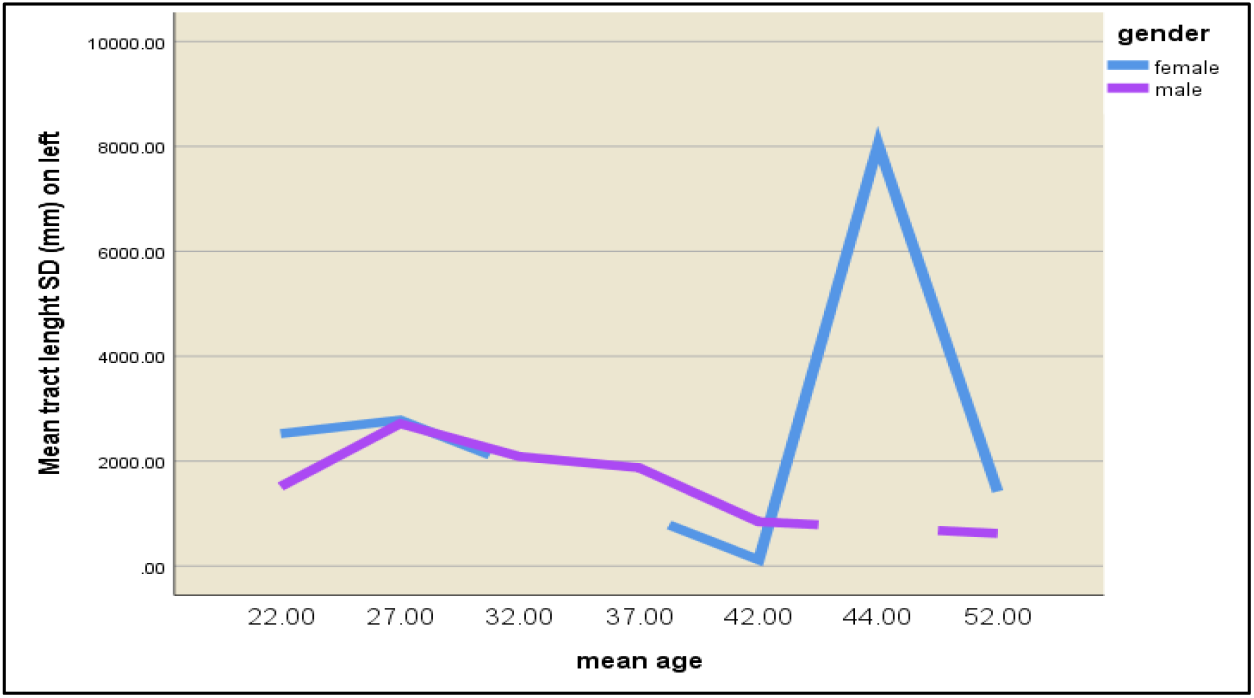
Graphical representation of tract length standard deviation on both right and left sides in male subjects.

We found that female subjects have a greater tract length standard deviation than in male subjects on left side.

## DISCUSSION AND CONCLUSION

The Dorsal (Occipitoparietal) pathway is a part of the dual stream visual brain theory where fibers span from the primary visual cortex to the parietal lobe. The dorsal stream is incapable of visual memory and is affiliated with “on line” control of actions and the spatial localization of stimuli and visual guidance of motor actions; “where?” and “how?” visuospatial discrimination [9]. With the use of Diffusion Tensor Magnetic Resonance Imaging Tractography, we were able to study the structural connections of the primary visual cortex (V1) to the Superior Parietal lobe in efforts to relate its clinical and functional significance [10]. “Skilled grasp can be defined as hand movements requiring independent control of each finger, which has been shown to rely on a highly developed corticospinal tract (<u>Lemon, 2008</u>)” [9]. When reaching out to grasp an object, the interpretation of sensory information is critical in accurately performing the action in terms of adaptation to the size and location of the object [11], [13], [15]. The object has to be visually distinguished before commencing the desired motion that would be executed into the motor action plan. Fibers of the occipitoparietal pathway projects to premotor areas PMv and PMd (Polanena.VDavare.M (2015) allows for visual guidance in the performance of skilled hand movements [9]. As such, lesions of the fibers in the dorsal stream (Posterior Parietal Cortex) would negatively influence spatial perplexity of individuals characterized by a deficit in the acuity of relative loci and the localization of objects during targeted actions resulting in Optic ataxia [11]. Optic ataxia is a neurodegenerative disorder that is prevalent in patients suffering from Alzheimer’s disease (Posterior cortical atrophy-the visual variant of Alzheimer’s disease) in which there is a higher order visual dysfunction. Optic ataxia is a part of the triad of neurophysiological impairments in Balint’s Syndrome in which there is a loss of the ability to coordinate manual movements [11], [12], [14]. In relation, lesions to the Posterior Parietal Cortex (PPC) would result in patients having a loss of the ability to coordinate manual movements in reference to their peripheral visual camp hence development of Optic Ataxia due to the inability to be visually guided by the dorsal stream visual pathway [13]. Patients therefore would not only encounter difficulty in performing low resolution spatially accurate reaches, but would also experience challenges when trying to adjust their hand posture in reaching out to grasp objects of varying alignment or size [15]. For example, in a case that a patient has a lesion in parietal posterior cingulate (lesion before the optic decussation), that patient would think that his hand is loose but he does not realize that he cannot coordinate his fine hand movements beyond the nasal camp on the affected visual site. We now conclude the findings of this study by stating; the current observations propose further insights to understand the structural existence and functional correlations for visuo-motor coordination pathway or “how” stream pathways in visual perception. Damage to this “how “stream fibers in the Visual pathways, manifest as Optic ataxia in Alzheimer’s Patients. However, these findings need to be confirmed with functional MRIs analysis in future understandings for better understanding of its significance in other neurodegenerative diseases and how this finding can affect future diagnostic and treatment approaches in these disorders.

